# The diverse effects of phenotypic dominance on hybrid fitness

**DOI:** 10.1101/2021.06.30.450598

**Authors:** Hilde Schneemann, Aslı D. Munzur, Ken A. Thompson, John J. Welch

## Abstract

When divergent populations interbreed, their alleles are brought together in hybrids. These hybrids may express novel phenotypes, not previously exposed to selection. In the initial F1 cross, most divergent alleles are present as heterozygotes. Therefore, F1 fitness can be influenced by dominance effects that first appear together in the hybrids, and so could not have been selected to function well together. We present a systematic study of these F1 dominance effects by introducing variable phenotypic dominance into Fisher’s geometric model. We show that dominance often reduces hybrid fitness, which can lead to patterns of optimal outbreeding and a steady decline in F1 fitness at high levels of divergence. We also show that “lucky” beneficial effects sometimes arise by chance, which might be especially important when hybrids can access novel environments. We then explore the interaction of phenotypic dominance with uniparental inheritance, showing that dominance can lead to violations of Haldane’s Rule (reduced fitness of the heterogametic sex) while strengthening Darwin’s Corollary (fitness differences between cross directions). Taken together, our results show that dominance could play an important role in the outcomes of hybridisation after secondary contact, and thus to the maintenance or collapse of isolating barriers. Nevertheless, the telltale signs of dominance are relatively few and subtle. Results also suggest that dominance effects are smaller than the cost of segregation variance, implying that simple additive models may still give good predictions for later-generation recombinant hybrids, even when dominance qualitatively alters outcomes for the F1.

## Introduction

During hybridisation alleles from diverged genomes can be expressed together for the first time. The interactions between these alleles will help to determine the outcome of the hybridisation. If hybrids are sufficiently fit, for example, then the hybridisation might lead to ongoing gene flow (e.g. Rieseberg et al., 1999; Lee et al., 2013), or to a hybrid swarm, potentially adapted to a new niche (e.g. Taylor et al., 2006). If hybrids are are unfit, by contrast, the lineages will remain reproductively isolated, and may be subject to reinforcement selection. All hybrids trace their ancestry to a first-generation, or ‘F1’ hybrid, and so the fitness of the F1 has a special importance. Ongoing gene flow, for example, is only possible if the F1 are both viable and fertile (Wallace, 1889; Fisher, 1930; Dobzhansky, 1937; Blair, 1955; Butlin, 1987). Moreover, the F1 are often easier to form than later generation crosses (Muller, 1940; Vetukhiv, 1954), and so have been more extensively studied.

Data from F1 hybrids show some recurring patterns, that have been observed consistently across taxa (Arnold and Hodges, 1995; Coyne and Orr, 2004; Fraïsse et al., 2016, Table S1). For example, crosses between closely-related parental lines sometimes exhibit “optimal outbreeding”, where the F1 are fittest between parents of intermediate genetic distance (Bateson, 1978; Butlin, 1987; Waser, 1993; Wei and Zhang, 2018; Dagilis et al., 2019); while at higher divergences, data often show an “F1 clock”, where F1 fitness declines steadily with genetic distance (Edmands, 2002; Coughlan and Matute, 2020). Perhaps best known is Haldane’s Rule, which describes the fitness inferiority of heterogametic F1 (Haldane, 1922; Coyne and Orr, 2004, Ch. 8; Schilthuizen et al., 2011); but equally well supported is “Darwin’s Corollary”, which describes strong fitness differences between the reciprocal F1 (i.e. female-male vs. male-female cross directions of the same parental lines; Darwin, 1859, Ch. 8; Kölreuters, 1766; Turelli and Moyle, 2007).

A class of fitness landscapes based on Fisher’s geometric model (Fisher, 1930; Orr, 1998) has been successful in generating many of these empirical patterns (Mani and Clarke, 1990; Barton, 2001; Chevin et al., 2014; Fraïsse et al., 2016; Simon et al., 2018; Schneemann et al., 2020). However, the model makes some unrealistic predictions for the F1; in particular the simplest form of the model predicts F1 heterosis (i.e., hybrid fitness advantage) at all levels of genetic divergence (Barton, 2001; Fraïsse et al., 2016). One explanation for this incorrect prediction is that most studies of Fisher’s model assume phenotypic additivity—that is, they assume that the phenotypic effect of a homozygous allele is exactly double its heterozygous effect. By contrast, empirical studies indicate that variable levels of dominance are common for loci affecting quantitative traits, including fitness components (Lynch and Walsh, 1998, p.485). Given its high level of heterozygosity, the F1 will be especially affected by the effects of this variable dominance.

Here, building on previous work (Schneemann et al., 2020), we present a systematic exploration of the effects of variable phenotypic dominance on hybrid fitness, with a particular focus on the F1. We show that the effects of dominance are highly diverse, but can nonetheless be divided into two broad categories: predictable deleterious effects, and “lucky” beneficial effects, which occur only occasionally and unpredictably. We then explore the interactions of dominance with uniparental inheritance or expression (as is essential to Haldane’s Rule and Darwin’s Corollary). We show that the two classes of effect act in similar ways, but that their interactions are sometimes synergistic and sometimes antagonistic. We end by discussing the overall importance of dominance to the outcome of hybridization.

## Model

Fisher’s (1930) geometric model assigns fitnesses to genotypes using a simple model of *n* quantitative traits under optimizing selection. The parameter *n*, which is sometimes called “organismal complexity”, is also the dimensionality of the fitness landscape. An individual’s phenotype is represented by a point in this landscape, with its trait values collected in the *n*-dimensional vector, **z**. The relative fitness of the individual is denoted as *w*, and depends on the Euclidean distance of the phenotype from some optimum **o**, whose position is determined by the current environmental conditions. Here, we will use a simple quadratic model, where log fitness declines with the squared distance to the optimum:

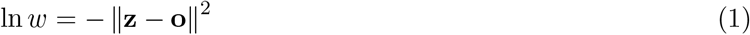

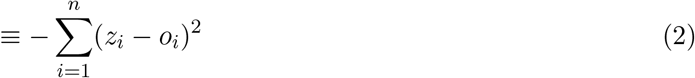

Substitutions are modelled as *n*-dimensional vectors of change to the phenotype. Thus, any two phenotypes can be connected by a chain of vectors that represent the divergent alleles accrued since their most recent common ancestor. This is illustrated in Figure 1a, where we show a chain of *d* = 6 genomic differences connecting two homozygous parental lines, labelled P1 and P2. In the illustration, each population fixed equal numbers of substitutions, but this is not an assumption of the analyses. What is important, however, is that all substitutions are defined relative to the P1 genotype. This means that none of our results depend on knowing whether the P1 alleles are ancestral or derived.

**Figure 1:**
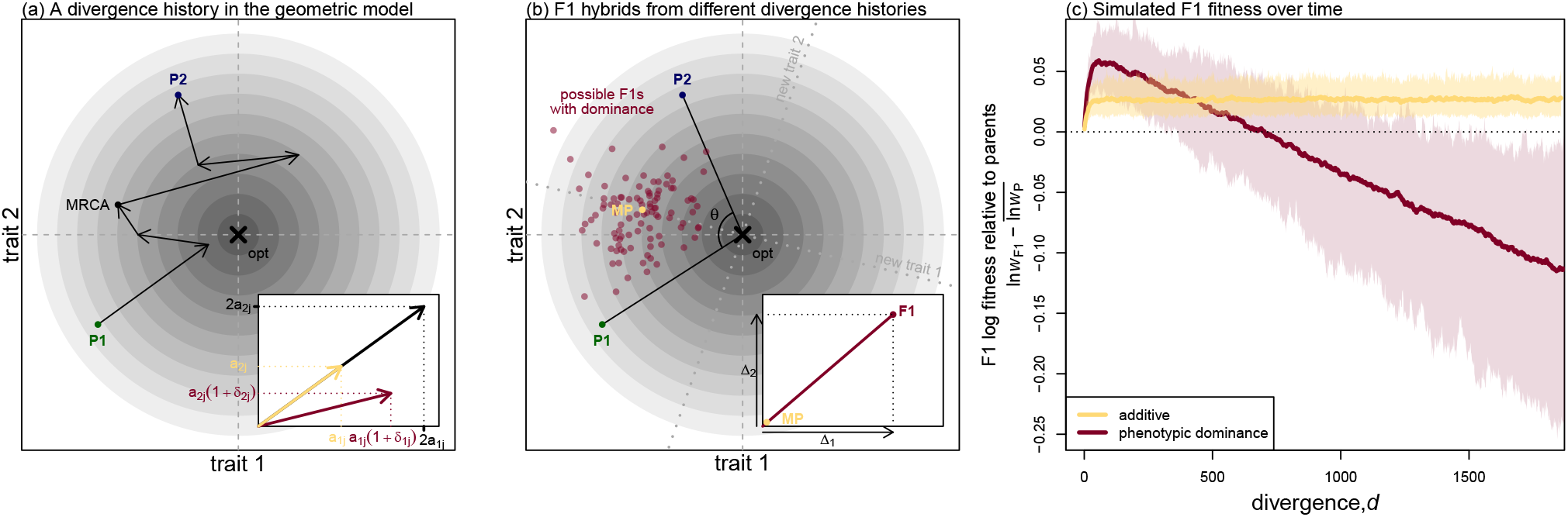
The effects of variable phenotypic dominance on F1 hybrid phenotypes and fitness in Fisher’s geometric model. **(a)**-**(b)**: illustrations of the fitness landscape model with *n* = 2 traits for visualisation. Contour lines indicate fitness, with darker colours closer to the optimum (× symbol). **(a)**: a pair of parental lines, P1 and P2, connected by a chain of *d* = 6 fixed differences. These pass through their most recent common ancestor (MRCA), but are orientated from P1 to P2. The inset shows a single substitution in homozygous state (black arrow), and in heterozygous state either with or without dominance. With additivity (yellow arrow), the heterozygous substitution is simply half of the homozygous substitution. With dominance (dark red arrow), the heterozygous effect on trait *i* is *a*_*ij*_(1 + *δ*_*ij*_), where *a*_*ij*_ is the additive effect, and *δ*_*ij*_ the dominance coefficient. **(b)**: parents and possible F1 hybrids. The yellow point (MP) shows the midparental phenotype located halfway between the parents, which is equivalent to the F1 under strictly biparental inheritance and additivity. Phenotypic dominance pushes the F1 phenotype away from the midparent. Dark red points show a cloud of possible F1 that could have resulted from different realisations of the divergence history. The inset panel shows the total dominance deviation on each trait (Δ*_i_*, eq. 4), and dotted lines indicate the new axes after the rotation implied by eq. 5. **(c)**: simulation results illustrating the difference between F1 and parental log fitness over the course of divergence. Under additivity (yellow lines), the F1 maintains a roughly constant level of heterosis. With phenotypic dominance (dark red lines) the F1 shows a pattern of optimal outbreeding with an initial spike in F1 fitness followed by a steady decline (the F1 clock). Lines show the means across 100 replicate simulations and the shading shows 95% quantiles. The simulation procedure assumed that larger effect mutations were more likely to be recessive (Fig. S1), and is described in full in Appendix 2. Parameters were *n* = 20 and 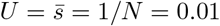.

Any given hybrid between P1 and P2 will contain some combination of the *d* vectors. Under phenotypic additivity, the effects of these vectors sum together both between and within loci. This means that an allele carried as a heterozygote will have the same orientation, but half the length, of the allele carried as a homozygote. This is shown by the yellow arrow in Figure 1a inset. To model phenotypic dominance, we relax the assumption of additivity within loci, using dominance coefficients *δ*_*ij*_. This is shown by the dark red arrow in Fig. 1a inset. Non-zero values of the *δ*_*ij*_ can alter both the length of the vector, and its orientation if different traits show different levels of dominance.

We note that the very simple model described above can be derived, either exactly or approximately, from a large family of more complex fitness functions (Martin and Lenormand, 2006; Martin, 2014). In that case, few if any of the *n* traits correspond to real quantitative traits, so that *n* becomes a phenomenological parameter of the fitness landscape (Orr, 2000; Welch and Waxman, 2003; Martin and Lenormand, 2006).

## Results

To explore the various effects of dominance on the fitness of F1 hybrids, we consider several different scenarios and patterns. As such, throughout the results section, we present the major results without derivation, and relegate the full derivations to Appendix 1. Simulations are used solely to illustrate the analytical results.

### 1. The F1 under additivity

With standard biparental inheritance and expression, F1 hybrids contain all *d* of the divergent alleles as heterozygotes. With complete phenotypic additivity, the F1 phenotype therefore matches the midparental phenotype, obtained by averaging the parental values for all *n* traits. The midparent is illustrated by the yellow point labelled MP in Figure 1b.

The fitness of the midparental phenotype, *w*_MP_, therefore depends solely on the fitnesses of the parental lines, *w*_P1_ and *w*_P2_, and their relative positions in the *n*-dimensional phenotypic space. This is characterized by *θ*, the angle in radians between the parental phenotypes (see Fig. 1b). Indeed, from the definition of cosine similarity, we find:

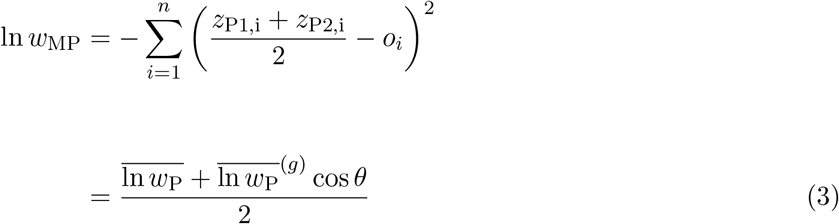

where 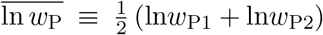 and 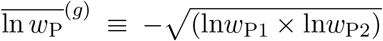 are respectively the arithmetic and geometric mean log fitness of the parents. Since cos *θ* varies between −1 and 1, it follows that 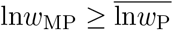. Therefore, with phenotypic additivity, the F1 will always be at least as fit as the average of the fitnesses of the two parental lines (Barton, 2001; Fraïsse et al., 2016).

To understand the consequences of this result, let us consider a concrete situation, in which the two parental lines remain well adapted to a fixed phenotypic optimum but diverge from their common ancestor via “system drift” (Schneemann et al., 2020; Schiffman and Ralph, 2021). In the initial stages of divergence the parental lines fix different alleles, and so they wander slightly in phenotypic space while remaining near the optimum. This wandering tends to increase *θ*, bringing the midparent closer to the optimum and thereby increasing the potential for heterosis (eq. 3). After this initial divergence, heterosis remains at roughly constant levels because the phenotypic distance has no further tendency to increase. This is confirmed by individual-based simulations of Fisher’s geometric model, which are described in detail in Appendix 2 and shown by the yellow lines in Figure 1c. With a fixed optimum and additive phenotypes, F1 heterosis shows a brief initial increase before settling at a constant positive value. The prediction of heterosis at all levels of genetic divergence is perhaps the least realistic prediction of the additive model.

### 2. The deleterious effects of phenotypic dominance

The principle effect of variable phenotypic dominance is to push the F1 phenotype away from the midparent, usually reducing its fitness (Schneemann et al., 2020). To see this, let us first define the total dominance deviation on trait *i*, including contributions from all *d* substitutions.

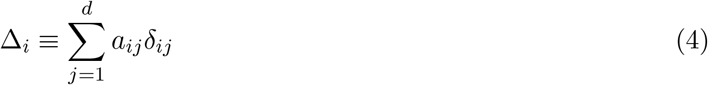

Here, *a*_*ij*_ is the additive effect of substitution *j* on trait *i*, and *δ*_*ij*_ is its dominance coefficient. These quantities are illustrated in the inset panels in Figure 1a and 1b. With this definition, the fitness of the F1 can be written as:

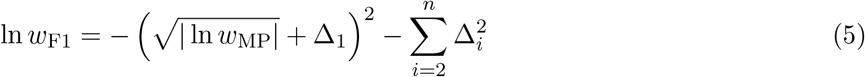

Here, without loss of generality, we have used a simple rotation of the trait axes such that the midparent falls short of the optimum only for trait 1 (as newly defined), but is optimal for the remaining *n* − 1 traits. This is shown in Figure 1b.

An important point about eq. 5 is that the Δ*_i_* affect fitness only in hybrids, and so unlike the additive effects, which act together in the P2 genotype, the dominance effects are unlikely to be under selection together during divergence, especially allopatric divergence. As such, they show no tendency to be coadapted, or remain close to any optimum. This means that similar scenarios of parental divergence, with similar trajectories for the parental phenotypes, can nonetheless yield very different Δ*_i_* and thus very different F1 phenotypes. This is illustrated by the cloud of dark red points in Figure 1b, which show possible F1 that might have resulted from the same divergence scenario between P1 and P2.

To predict F1 fitness, we must average over these possible evolutionary histories. Let us first consider the scenario discussed above, where parental populations remain well adapted to the same constant environment. In this case, we can think of the Δ*_i_* as undergoing a random walk away from the midparent, so that the set of possible F1 form a growing cloud (Schneemann et al., 2020). This yields:

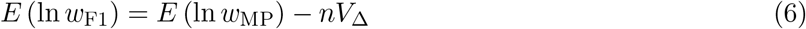

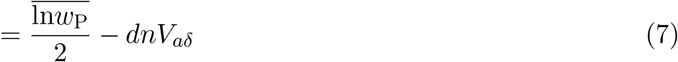

where *V*_Δ_ is the variance, across divergence histories, of the the total dominance deviations (see eqs. 4 and 25); while *V*_*aδ*_ is the equivalent variance in dominance effects of single substitutions (i.e. the *d* increments of the random walk; eq. 26). As with the additive model, eqs. 6–7 predict a transient increase in F1 fitness in the early stages of divergence, as long as the second terms remain small. But as the number of substitutions (*d*) increases, the second terms start to grow, causing a steady fitness decline. To illustrate this, we repeated our simulations after allowing for variable phenotypic dominance among the new mutations (see Appendix 2 for full details). The results, shown as dark red curves in Figure 1c, show a clear pattern of transient heterosis (i.e., optimal outbreeding) followed by a linear decline in log fitness (the F1 clock). The maximal heterosis is also higher with dominance, due to the parental lines fixing more deleterious recessives.

We can also think of the results above as exemplifying two regimes. In the early stages of divergence, the phenotypic distance between the parents increases faster than the cloud of possible F1. This means that most F1 trait values will be intermediate between the parental values. Later, the parents stop diverging phenotypically (because of effective stabilizing selection acting on their traits), but the cloud of possible F1 continues to expand. This leads to transgressive trait variation, with the F1 values lying outside of the range of the parental values. It is notable that both of these regimes have been observed in quantitative traits from F1 hybrids: transgressive variation sometimes decreases with *d*, and sometimes increases with *d* (Stelkens and Seehausen, 2009, see also Appendix 1 and Figure S2 for more details).

#### 2.1. The tick rate of the F1 clock

Equation 7 gives the tick rate of the F1 clock as *nV*_*aδ*_, which is also the rate of expansion of the cloud of possible F1 phenotypes. This quantity summarizes the distribution of factors fixed, which, as previous work has shown, will depend on many different aspects of the system’s biology (e.g. Charlesworth, 1992; Orr, 1998; Griswold, 2006; Yeaman and Whitlock, 2011; Matuszewski et al., 2014; Schneemann et al., 2020; Yamaguchi and Otto, 2020).

One important determinant of the tick rate is likely to be the distribution of dominance coefficients of new mutations. This is easiest to see with a simpler model of mutation than that used in Fig. 1c. If we have isotropic traits i.e. equal distributions of effects across traits, and no tendency for substitutions with larger additive effects to have more extreme dominance coefficients (e.g. no covariance between 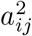 and 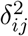), then we have

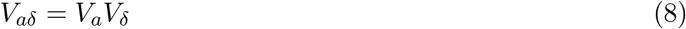

where *V*_*a*_ and *V*_*δ*_ are the variance of the additive effects and dominance coefficients respectively. Further-178 more, with this model, *V*_*δ*_ for the fixed effects will often match the equivalent quantity for new mutations, denoted 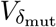. To see this, we repeated our simulations with the assumptions just described. The dominance coefficient of each new mutation on each trait was drawn from a shifted beta distribution, with a vanishing mean (such that mutations were semi-dominant on average) and variance 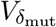. We simulated under seven different variances, including the extremes of 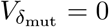 (phenotypic additivity) and 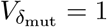, such that new mutations were either fully dominant or fully recessive with a 50:50 probability. The seven distributions we used are illustrated in Figure 2a.

**Figure 2:**
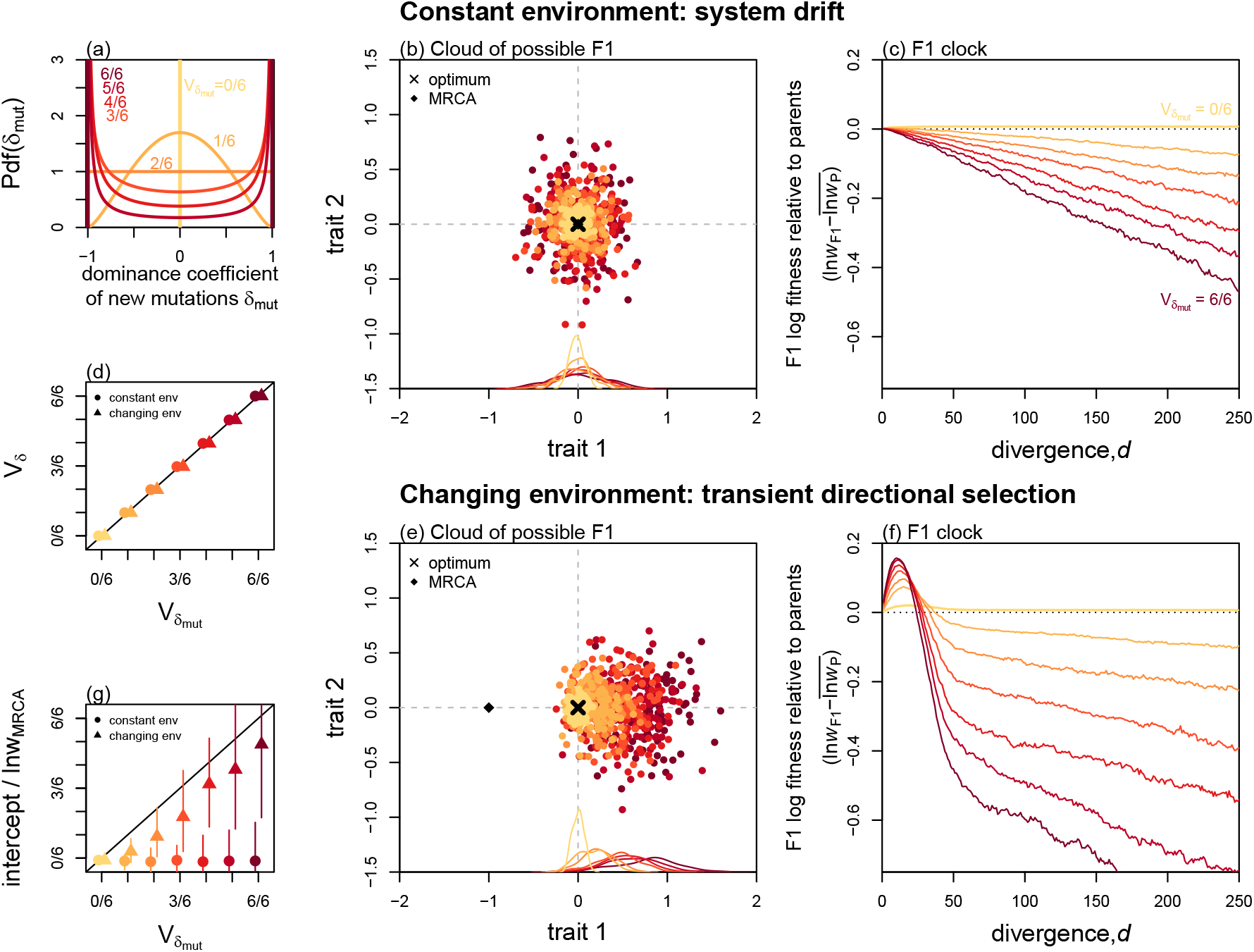
Deleterious effects of phenotypic dominance depend on the distribution of new mutations and the history of directional selection. **(a)**: The distribution of dominance coefficients for new mutations used in each set of simulations. These were 7 different beta distributions, with vanishing means and 7 different variances, ranging from additivity (yellow: 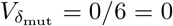) to complete dominance or recessivity (dark red: 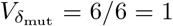). **(b)**–**(c)**: Simulations under constant environmental conditions show that **(b)** the variance in the cloud of possible F1s, and the tick rate of the F1 clock both increase steadily with 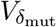. **(d)**: This is because, for these simulations, the dominance coefficients of fixed mutations reflect those of the new mutations, such that 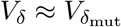. **(e)**: Directional selection can lead to directional dominance, causing the cloud of possible F1s to ‘overshoot’ the optimum in the direction of the past adaptation. **(f)**: Directional dominance creates transient heterosis as the parental lines evolve towards the new optimum, but a deleterious overshoot once the new optimum is reached. **(g)**: The result is that, after an initial period, the F1 clock ticks at the same rate, but with an intercept which also increases with 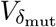. (b) and (e) show 100 simulations stopped after *d* = 50 substitutions, and with parameters chosen to best visualise the clouds (*N* = 100, *n* = 2). (c)–(d), and (f)–(g) show means across 100 replicate simulations with *n* = 20 and *N* = 1000 to generate a steady clock. All panels use 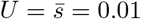.

Figure 2b shows clouds of F1 phenotypes for sets of 100 replicate simulations, each with *n* = 2 for visualisation. In all cases, the cloud of possible F1 is centred on the optimum, but grows in width with 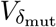. Figure 2c shows the consequence of this F1 cloud expansion for the F1 clock. Here, to match Fig. 1c, we simulated with *n* = 20 traits, but with a larger population size, so that parental populations remained well adapted and thus had little potential for F1 heterosis. Results show that the F1 tick rate also increases steadily with the variance in mutational dominance, 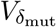. Figure 2d (circles) shows that this is because, as anticipated, the dominance coefficients of the fixed substitutions closely match the dominance coefficients of new mutations (i.e., 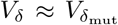; see also Schneemann et al. 2020). As shown in Figure S3, 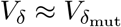 does not hold in all parameter regimes. However, the increase in the slope with 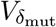 was consistently observed, including when each value of *δ*_mut_ applied to all traits, so that heterozygous and homozygous mutations differed in size, but not direction.

#### 2.2. Directional selection and directional dominance

Results above assume that dominance deviations have no tendency to point in any phenotypic direction, such that the cloud of possible F1s is centered on the midparent, and all *E* (Δ*_i_*) = 0 (see Fig 1b). However, it is well known that directional selection can lead to directional dominance, as with Haldane’s Sieve (Haldane, 1924, 1927; Frankham, 1990; Crnokrak and Roff, 1995; Ala-Honkola et al., 2013; Sztepanacz and Blows, 2015; Kapun et al., 2016). This is because populations evolving in a particular direction preferentially fix alleles that are dominant in that direction.

To understand the effects of directional dominance for F1 fitness, let us consider the most extreme case, where both populations adapt independently to identical environmental change, so that all of the dominance deviations point in the same direction. In this case, the cloud of possible F1 becomes shifted in the direction of the past evolutionary change. This is illustrated in Figure 2e, where a bout of directional selection on trait 1 leads to *E*(Δ_1_) > 0. To understand why this happens, let us consider the evolution of trait 1 during the bout of adaptation. If we denote the F1 trait as *z*_F1,1_, and the midparental value as *z*_MP,1_, then eqs. 2 and 5 imply that:

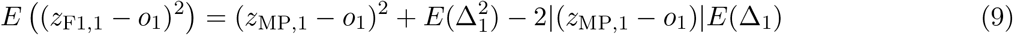

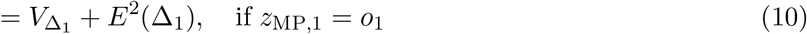

The important point about eq. 9 is that it contains both positive and negative terms. This is because, during the adaptation phase, the directional dominance can increase F1 fitness, by taking its trait value closer to the still-distant optimum. But as shown by eq. 10, once the new optimum is approached, the directional dominance leads to a permanent and deleterious ‘overshoot’ of this optimum (see also Figure S4, and Ono et al. (2017) for a related phenomenon shown in yeast).

If the adaptation to the new optimum required *d*_dir_ substitutions, the expected log fitness of the F1 after this period (i.e. once system drift at the new optimum has begun) is:

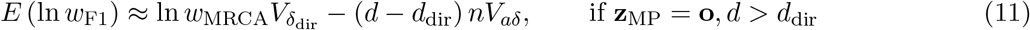

where 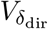 is the variance in dominance coefficients of the adaptive substitutions, and ln *w*_MRCA_ is the log fitness of the maladapted ancestor. Comparing eq. 11 to earlier results (eqs. 6 and 7), we see that the overshoot adds a new term. The history of directional selection leads to a non-zero intercept for the F1 clock.

Complete simulations of this scenario are reported in Figure 2f. Comparison of Figures 2f and 2c shows clearly the transient increase in F1 fitness (eq. 9), and the deleterious overshoot once the new optimum is approached (eq. 10). After the initial period, the F1 clock continues to tick at the same rate as before (see triangles in Figure 2d), but with a permanent intercept (eq. 11). Fitting linear regressions to the F1 clocks in Figure 2f after excluding the first 50 substitutions allowed us to calculate this intercept. Figure 2g (triangles) confirms that the intercept is also affected by the input of new mutations, such that 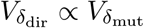.

Results above concern directional selection in a common direction, but they generalize readily. For example, if each parental population underwent directional adaptive change on a different phenotypic trait, the resulting directional dominance would lead to an F1 with a mixture of the derived traits of the two parental lines. This is the case described as trait mismatch by Thompson et al. (2021). In all cases, with variable phenotypic dominance, a history of past directional selection can lead to an additional loss of fitness for the F1.

### 3. The lucky beneficial effects of dominance

So far, we have considered the typical effects of dominance in the F1, by averaging over the possible evolutionary histories. These effects are generally deleterious, and any heterosis tends to be transient. Nevertheless, even when effects are deleterious on average, by chance alone, some realisations of the evolutionary process will take the F1 closer to the optimum. In Fisher’s geometric model, these “lucky” outcomes are far more likely when the number of phenotypic traits, *n*, is small. This is because random changes are more likely to go in the right direction when the dimensionality of the landscape is low (Fisher, 1930; Orr, 2000).

To see this, let us first consider again the scenario illustrated in Figures 1c and 2b–c, where parental populations remain well adapted to a single fixed optimum. We assume that the Δ*_i_* have vanishing means (i.e., no directional dominance), and are approximately normally distributed. The normality is justified by the central-limit-like behaviour arising from the sum in eq. 4. With these assumptions, the coefficient of variation in log F1 fitness is approximately

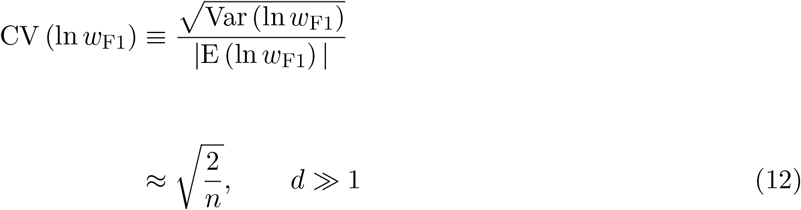

Equation 12 implies that, when *n* is large, the F1 clock will tick in a relatively deterministic way. When *n* is small, by contrast, increasing the divergence between the parental lines may lead to the chance re-appearance of heterosis, even after many generations of low fitness F1. This is confirmed by simulations reported in Figure S5a–b.

#### 3.1. Lucky beneficial effects in a novel environment

The lucky beneficial effects of dominance may be particularly consequential when hybrids are formed in novel environments, to which one or both parental lines are severely maladapted. In the additive model, F1 heterosis can only appear when the parents are maladapted in different ways (Simon et al., 2018; Schneemann et al., 2020; Yamaguchi and Otto, 2020). For example, hybrids between parents adapted to low and high altitudes might thrive at intermediate altitudes (Wang et al., 1997). With phenoptypic dominance, however, hybrid advantage might appear under a broader range of conditions. Consider, for example, the situation shown in Figure 3a, where two genomically divergent, but phenotypically similar parental lines hybridize in a novel habitat. In this case, the midparent matches the parental phenotypes (ln *w*_MP_ ≈ ln *w*_P_), and so the effects of dominance will be deleterious on average (eq. 6). Nevertheless, for some divergence histories, the F1 will be fitter by chance. The probability of this lucky heterosis can be derived by noting that eq. 5 has a non-central chi-squared distribution if the Δ*_i_* are normal. The probability therefore depends on *n* and the ratio *V*_Δ_/ ln *w*_P_, which compares the sizes of the dominance deviations to the maladaptation of the parents. Figure 3b plots this probability for a range of values, and we also have the approximation:

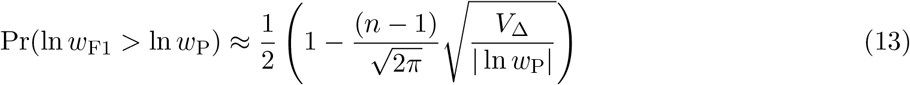

**Figure 3:**
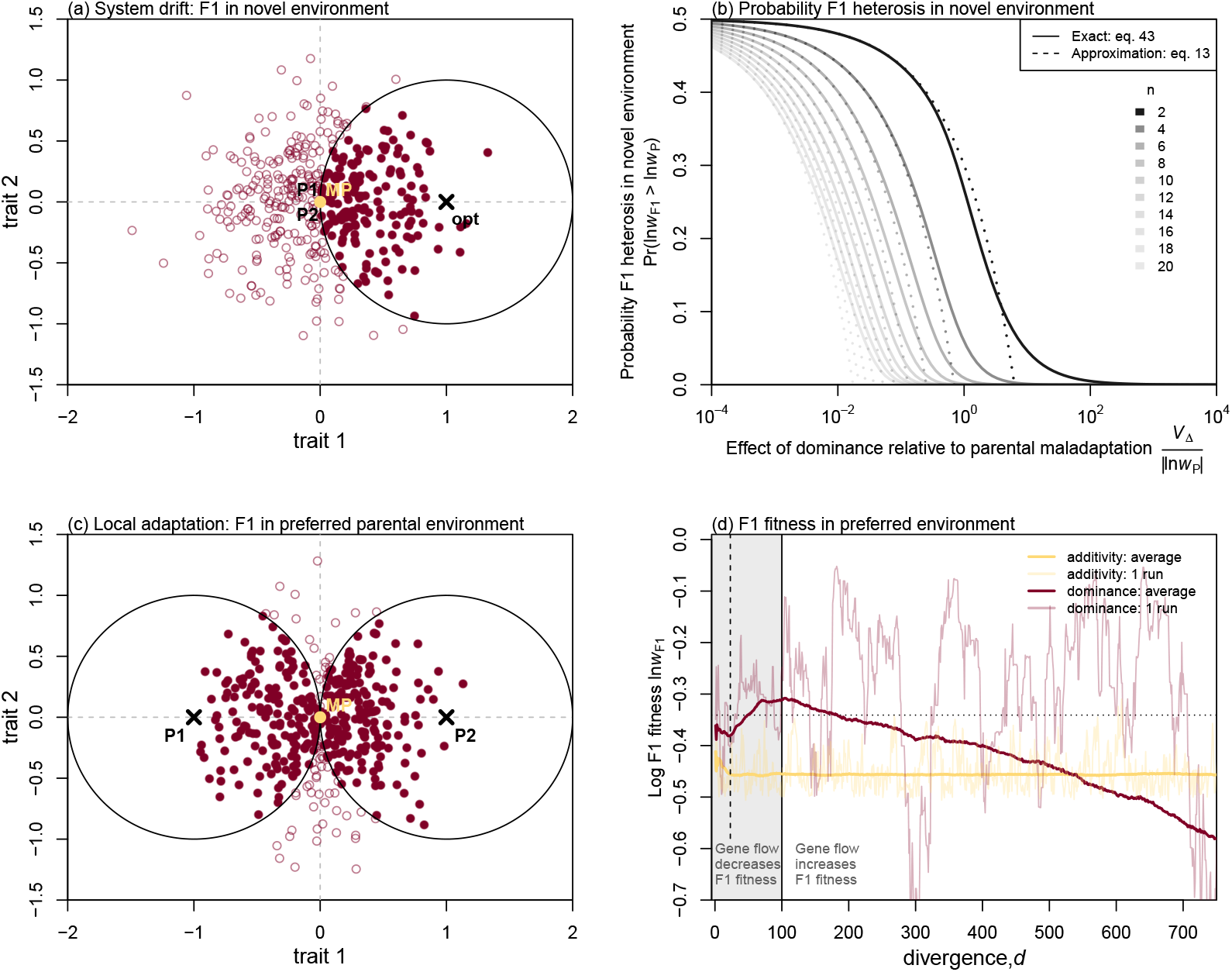
“Lucky” beneficial effects of dominance in heterogeneous environments. **(a)**: Even when two parental lines are well adapted to similar environments, their F1 may be well adapted to a novel environment, simply by chance. These fitter F1 (solid dark red points) lie within the circle shown, with the new optimum at its centre. **(b)**: The probability of such fitter F1 appearing decreases rapidly with *n* (the dimensionality) and with *V*_Δ_/| ln *w*_P_| (the size of the dominance deviations relative to the maladaptation of the parents). **(c)**: When the parents are locally adapted to two different environments, dominance might yield an F1 that is much fitter in one of the two environments, potentially leading to asymmetrical gene flow. **(d)**: Simulations of the F1 of locally adapted parents. F1 were scored in the “preferred” parental habitat, i.e. the habitat to which they were better adapted. Simulations began with the MRCA intermediate between the two optima, and the parents adapted to their optima rapidly, always before *d* = 25 (see vertical dotted line). With additivity, F1 fitness remained roughly constant thereafter (yellow curves), but with dominance, expected F1 fitness continued to increase with divergence (dark red curves), until a maximum roughly predicted by eq. 15 (horizontal dotted line). However, changes in F1 fitness were erratic for any single run (see fainter curves). The result is that gene flow, which reduces *d*, can either increase or decrease F1 fitness with dominance. Simulations in (d) were the same as in Fig. 2e, with the extremes of 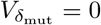 and 1, and averages taken over all possible pairs of the 200 parental populations; see also Figure S7.

(see Appendix 1). As shown in Figure 3b, the maximum probability of heterosis is 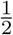, and it applies when dominance effects are small compared to the maladaptation of the parents such that *V*_Δ_ ≪ | ln *w*_P_| (toward the left of Fig. 3b). This implies that any fitness gain due to heterosis would also be very small, and simply reproduces Fisher’s result that very small changes have a 50% chance of being beneficial (Fisher, 1930). Conversely when the dominance effects are very large, such that *V*_Δ_ ≫ | ln *w*_P_| (toward the right of Fig. 3b), the F1 are almost certain to overshoot the new optimum. Therefore, the area of interest concerns *V*_Δ_ ≈ | ln *w*_P_| (toward the centre of Fig. 3b), and in this regime, the probability of heterosis declines rapidly with *n* (see also Fig. S6). Only when *n* is small is there a non-negligible chance that dominance effects are both substantial and beneficial.

#### 3.2. Hybrids between locally adapted parents

Now let us consider the situation shown in Figure 3c. This panel represents the outcome of divergent selection (Schluter, 2000), where each parental line is well adapted to a different habitat, characterized by different phenotypic optima. In this case, the midparent lies midway between the optima, such that, in either habitat ln *w*_MP_ ≈ ln *w*_P_/4, with ln *w*_P_ denoting the fitness of the maladapted parental line. In this scenario, results in Figure 3b now describe the probability of the F1 being fitter than the midparent in one of the two parental habitats. In most cases, the F1 will be fitter in one of the habitats than the other, and in this “preferred habitat”, its expected log fitness is:

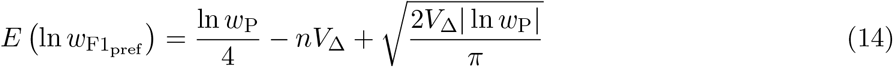

Equation 14 neatly separates the negative and positive effects of dominance in its second and third terms, and shows that the deleterious effects—but not the beneficial effects—grow with *n*. Equation 14 also shows that the fitness benefits of dominance will be greatest at intermediate values of *V*_Δ_, and therefore, at intermediate levels of divergence (eqs. 6–7). This means that, in contrast to the scenario shown in Fig. 2e–f, F1 fitness in one of the two habitats, might continue to increase even after two parental populations have adapted to their new optima. At this intermediate level of divergence, when the fitness benefit is greatest, we find:

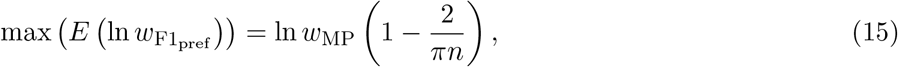

so, at suitable levels of divergence, phenotypic dominance can yield a substantial fitness increase for the F1 over the midparent. However, these lucky effects of dominance occur only in one of the two habitats and only when *n* is small. All of these results are confirmed in Figure 3d, which shows simulation results with *n* = 2 (see also Figure S7). Results confirm that on average (solid lines), F1 fitness with dominance initially increases with *d*, before reaching a maximum approximated by eq. 15, and then decreasing. However, eq. 12 still applies, and so for any single realisation of the divergence history (faint lines in Figure 3d) the fitness of F1 with phenotypic dominance is highly erratic.

Taken together, these results imply that phenotypic dominance may lead to asymmetrical gene flow between locally adapted populations after secondary contact. The asymmetry arises from fitness differences between the F1 in the two habitats and is therefore a form of “dominance drive” (Mallet and Barton, 1989; Barton, 1992). The results also imply that the gene flow, which by definition reduces *d*, may have an unpredictable effect on F1 fitness. In some cases, homogenisation of the parental genomes will increase the fitness of their F1, but in other cases, it will lead to a switch in the direction of the gene flow (as the F1 becomes adapted to the other parental habitat), or even to a substantially lower F1 fitness. In the latter case, the outcome would therefore resemble reinforcement selection, albeit via a completely different route.

### 4. The interaction of phenotypic dominance with uniparental inheritance

In addition to dominance, the F1 might differ from the midparent for a second reason: the uniparental inheritance or expression of divergent alleles, for example on sex chromosomes or mitochondria (Fraïsse et al., 2016; Simon et al., 2018). In the parental lines, the complete set of alleles may be co-adapted in their additive effects, e.g. due to compensatory changes. When hybrids inherit a complete set of parental alleles, these co-adapted relationships maintain intact; but when hybrids inherit some parts of their genome from only one parent, they may lack some of the co-adapted alleles from the other parent. As we show below, dominance and uniparental inheritance have similar effects on hybrid fitness, but with one key difference—under uniparental inheritance the direction of phenotypic deviation from the midparent is opposite for the two cross directions.

The effects of uniparental inheritance are illustrated in Figure 4a. In this figure, and throughout this section, we assume a concrete example of uniparental effects, although results generalize easily to other cases. Our example involves sex chromosomes, where the heterogametic sex are effectively XO (i.e. where males carry only the maternal X, and the Y is either missing or highly degenerate). We further assume a common form of dosage compensation where X-linked alleles have identical effects in homozygous and hemizygous state. This last assumption ensures that non-hybrid offspring of both sexes are phenotypically identical (i.e., that male and female offspring from a P1 × P1 cross are equally fit).

**Figure 4:**
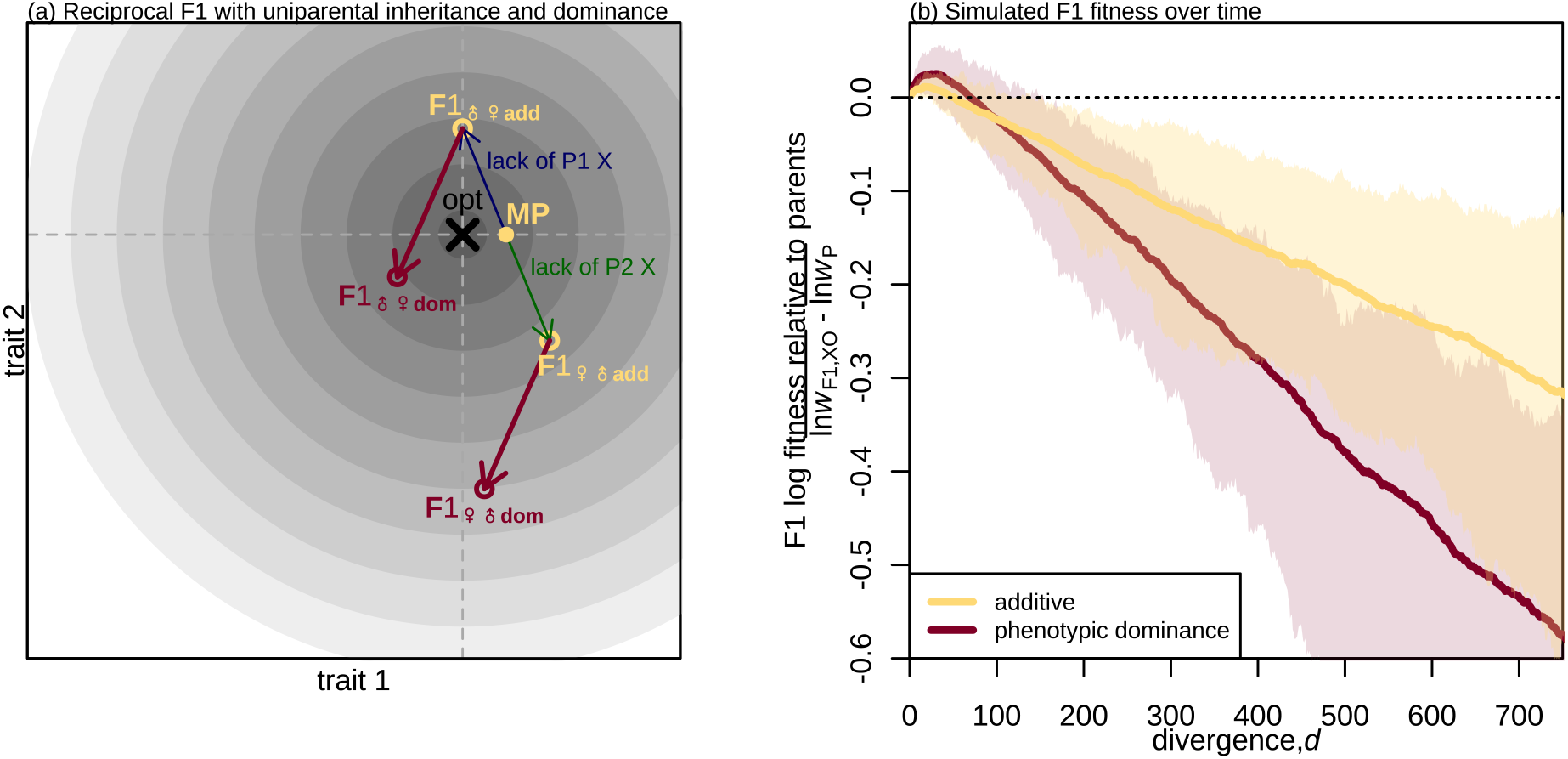
Uniparental effects interact with dominance to determine F1 fitness. Both phenotypic dominance and uniparental inheritance/expression cause the F1 to deviate from the midparent (MP). **(a)**: An illustration of heterogametic (XO) offspring, from a reciprocal F1 cross: i.e. including both the male-female and female-male cross directions of the same parentallines. Yellow points show the reciprocal F1 under additivity (F1_♀♂,add_, F1_♂♀,add_) and dark red points show the reciprocal F1 with phenotypic dominance (F1_♂♀,dom_, F1_♂♀,dom_). The dark blue and green lines indicate the deviations due to the absent paternal X, and are equal and opposite for the two cross directions. The dark red lines indicate the dominance deviations due to heterozygosity on the autosomes, and are identical for both cross directions. Both sets of deviations are expected to grow with the divergence, *d*. **(b)**: Simulation results of log fitness in XO F1 hybrids, averaged across the two cross directions. Results show that the F1 clock appears under additivity with uniparental effects. Plotting conventions and simulation runs are identical to those shown in Figure 1c, except that a proportion *x* = 1/4 of the divergent sites were randomly assigned to the X-chromosome before forming the hybrids. As a result, optimal outbreeding and the F1 clock appear not only with dominance (dark red curves) but also with additivity (yellow curves)

In the case described, the absence of the paternal X causes the heterogametic F1 to deviate from the midparent, even under phenotypic additivity. These deviations are equal and opposite for the two cross directions (see the blue and green arrows in Fig. 4a). If there is phenotypic dominance for alleles on the autosomes, then this leads to further deviations that apply identically to both cross directions (see the dark red arrows in Fig. 4a). In the sections below, we show how uniparental effects and phenotypic dominance combine to determine F1 fitness.

#### 4.1. The F1 clock under dominance and uniparental inheritance

If we consider any single F1, then uniparental effects and phenotypic dominance have essentially the same consequences. As such, they represent alternative, but non-exclusive explanations of the F1 clock and optimal outbreeding. To see this, let us consider the expected log fitness of an XO F1 hybrid. If we assume that a fraction, *x*, of the *d* divergent alleles are X-linked, then from eq. 5 and published results (Fraïsse et al., 2016; Simon et al., 2018), we find:

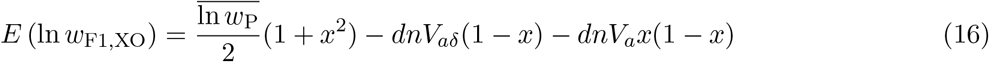

Equation 16 still predicts optimal outbreeding, but there are now two terms that reduce F1 fitness. The second term captures the dominance deviations from the autosomes, and the third term captures the lack of co-adaptation between paternal autosomes and the absent paternal X.

Figure 4b illustrates these results. We returned to our simulated data, and formed heterogametic F1 in the way described. Results show that optimal outbreeding and the F1 clock now appear under the additive model too (yellow curves; Fraïsse et al., 2016; see also Fig. S5c–d). Dominance simply accelerates the fitness decline (dark red curves).

#### 4.2. Haldane’s rule

If we consider fitness differences between heterogametic and homogametic hybrids, then the uniparental effects and phenotypic dominance tend to push in different directions. If the parents are well adapted, then the additive model predicts lower fitness for the heterogametic sex, in accord with Haldane’s Rule (Haldane, 1922; Barton, 2001; Fraïsse et al., 2016; Simon et al., 2018). This is due to the loss of co-adaptation, described above (see the third term of eq. 16). However, phenotypic dominance has the opposite effect, because it tends to makes heterozygosity deleterious, and heterogametic F1, being hemizygous for the X, lack the deleterious heterozygosity on the X (to see this, compare the second terms of eqs. 7 and 16).

To see how these two effects balance, let us consider the difference in log fitness between heterogametic and homogametic F1. Using eqs. 7, 8 and 16, we find:

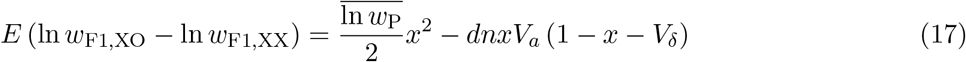

Haldane’s Rule holds on average if eq. 17 is negative. If parents are well adapted, or divergence is substantial, then the second term in eq. 17 dominates and we expect Haldane’s Rule on the condition that

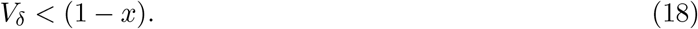

Note that this equation was derived under the restrictive assumptions of eq. 8, but it holds much more generally, if we define *V*_*δ*_ as the relative sizes of the dominance and additive effects (i.e., as *V*_*δ*_ ≡V_*aδ*_/*V*_*a*_, where the latter two quantities are variances averaged across traits).

Equation 18 confirms that Haldane’s Rule will always hold under additivity (when *V*_*δ*_ = 0), but that variable phenotypic dominance can lead to violations (i.e., ‘anti-Haldane’s Rule’), especially when the dominance coefficients are highly variable, or if the X is very large. This is confirmed by the simulations shown in Figure 5a–b. These plots use the simulation runs from Fig. 2b, for which the dominance coefficients match the mutational input 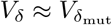. The results show that Haldane’s Rule is indeed violated when 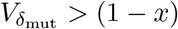.

**Figure 5:**
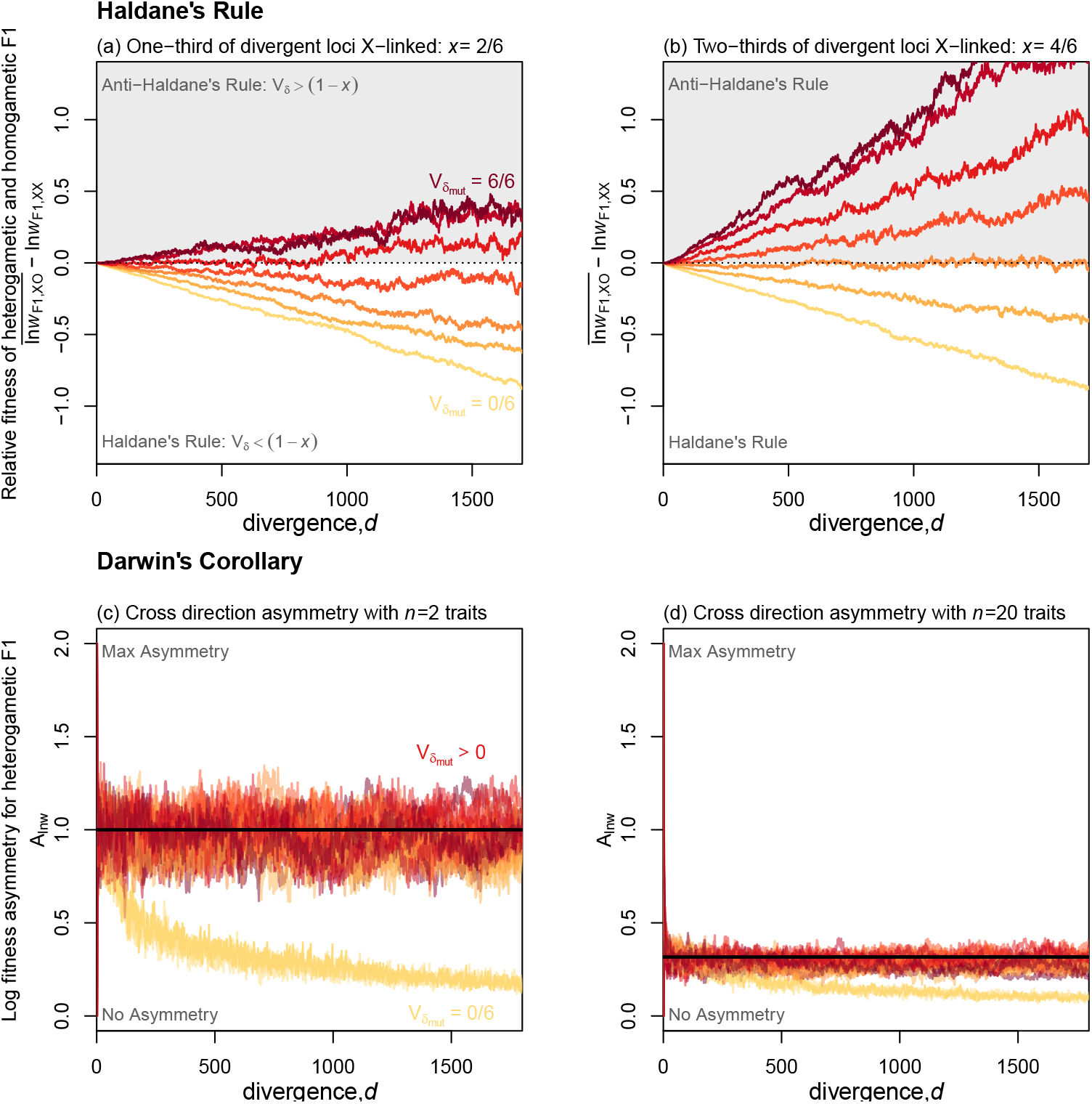
Differences between F1 cross types with uniparental inheritance and variable phenotypic dominance. **(a)–(b)**: Phenotypic dominance can lead to violations of Haldane’s Rule. Curves show the difference in mean log fitness between heterogametic and homogametic F1, at various levels of parental divergence. Curves below the horizontal dotted line indicate that heterogametic F1 are less fit, such that Haldane’s Rule holds. Colours correspond to those in Figure 2, and show the same 7 values of 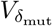. Haldane’s Rule always holds with additivity (yellow lines; 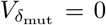), but can be violated if dominance coefficients are highly variable (darker colours), or if the X chromosome is very large – compare (a) to (b). **(c)–(d)**: Phenotypic dominance enhances log fitness asymmetry between reciprocal F1. Curves show the asymmetry measure of eq. 19, measured for XO hybrid offspring from the two cross directions. Results under additivity (yellow), differ qualitatively from those with phenotypic dominance (all other colours), and the latter all show approximately the same result: 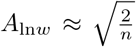 (black horizontal lines). Plots include 28 sets of simulations, using two populations sizes: *N* = 100, 1000; two sizes of X chromosome: 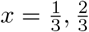; and the 7 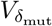 values shown in Fig. 2a). Each curve represents the mean across 100 simulation replicates, all with 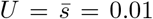; (a)–(b) used *n* = 20 and *N* = 100. Note that the divergence was simulated under strictly biparental inheritance, to ensure that 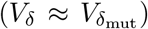.

#### 4.3. Darwin’s Corollary

Let us now consider the fitness differences between the F1 cross directions, that is a cross where P1 is dam (P1_♀_×P2_♂_) compared to a cross where P1 is sire (P2_♀_×P1_♂_). In stark contrast to Haldane’s Rule, phenotypic dominance works together with uniparental effects to generate this pattern. To see this, we will use the following measure of log fitness asymmetry, where the absolute difference between the cross directions is normalized by the absolute mean:

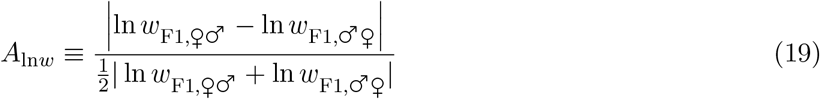

Here, ♀♂ and ♂♀ denote the two cross directions, and the statistic is bounded at 0 ≤ *A*_ln*w*_ ≤ 2.

Fraïsse et al. (2016) studied F1 asymmetry under the additive version of Fisher’s model, and showed that asymmetry could appear only if the midparental phenotype was maladapted. This reason is clear from Figure 4a. When the midparent matches the optimum, the equal and opposite deviations (blue and green arrows), will lead to identical fitness loss.

Furthermore, even if the midparent is maladapted, its distance from the optimum must be large relative to the deviations for substantial asymmetry to appear. But, in a stable environment, the parents (and therefore the midparent) remain close the optimum, while the deviations grow with *d* (e.g. eq. 7), such that the F1 in both cross directions are expected to become less fit, and the difference between their fitnesses smaller. This implies that any asymmetry between the cross directions will decline with *d* under the additive model. This is confirmed by the yellow curves in Figure 5c–d.

Adding phenotypic dominance to the model qualitatively changes this result. Because the dominance deviations also grow with *d*, levels of asymmetry can remain large, even at high divergence. An illustrative case is shown in Figure 4a. Moreover, with dominance, we find that to a rough approximation:

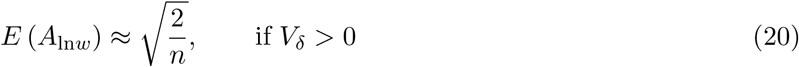

so the normalized asymmetry in log fitness is predicted to remain roughly constant at all levels of divergence, and to depend solely on the number of traits, *n*. This is confirmed by the simulations with dominance shown in Figure 5c–d. For all of the parameter values we simulated, as long as there was phenotypic dominance 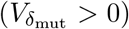, then asymmetry levels remained roughly constant, and were close to the prediction of eq. 20 (see also Figure S5e–f).

The arguments in this section show that under Fisher’s geometric model, Darwin’s Corollary can be explained as another “lucky” beneficial effect of dominance, where the good luck for one cross direction is balanced by bad luck for the other. The consequence is that – with phenotypic dominance, but not with additivity – Darwin’s Corollary is predicted at high levels of genomic divergence.

## Discussion

During evolutionary divergence between populations, the additive effects of the divergent alleles act together in homozygous genotypes, and so they will often by co-adapted – i.e. selected to function well together in the current environment. By contrast, the dominance effects will act together only in early generation hybrids, and so they are more likely to wander erratically in phenotypic space. We have presented a systematic study of these dominance effects and their consequences for F1 hybrid fitness. We have shown that the effects are usually deleterious, reducing hybrid fitness; but there can also be lucky outcomes when, by chance, the novel F1 phenotype is better adapted than the parental lines.

Taking the negative and positive effects together, dominance could have a broad range of consequences for the F1. These consequences include asymmetric and fluctuating gene flow between locally adapted populations (Fig. 3c–d), and F1 hybrid advantage, either in novel habitats (Fig. 3a–b), or during ongoing directional selection (Fig. 2f). The consequences also include well-known patterns, that have been observed in multiple taxa, such as optimal outbreeding and the F1 clock (Waser, 1993; Edmands, 2002; Wei and Zhang, 2018; Dagilis et al., 2019; Coughlan and Matute, 2020; Figs. 1c and 2c).

But while dominance is a possible cause of these patterns, it need not be their actual or only cause. Variable phenotypic dominance has often been measured directly (Lynch and Walsh, 1998, p.485; Thompson et al., 2021); including as transgressive trait variation, which is non-additive on any scale of measurement, and so cannot be easily “transformed away” (Lynch and Walsh, 1998, p.308). However, it does not follow that this dominance is important for determining hybrid fitness. While explicit links between traits and fitness can be made (e.g. Thompson et al., 2021), often the relevant set of traits and the way selection perceives these traits is unknown. In general, the “traits” in Fisher’s model need not be equated with standard quantitative traits (Martin, 2014), and an additive model may yield good predictions even if the measurable quantitative traits are highly non-additive.

Moreover, we have shown that some of the empirical patterns generated by dominance also appear under an additive phenotypic model, if some determinants of the phenotype are uniparentally inherited or expressed. Data give strong support for such uniparental effects – not least in the ubiquity of Darwin’s Corollary (Kölreuters, 1766; Darwin, 1859, Ch. 8; Muller, 1942; Turelli and Moyle, 2007), in diverse plant (Tiffin et al., 2001), fungus (Dettman et al., 2003), and animal systems (Bolnick et al., 2008; Brandvain et al., 2014), including simultaneous hermaphrodites (Escobar et al., 2008; Sato et al., 2014; Bouchemousse et al., 2016; Fraïsse et al., 2016).

One obvious question, therefore, is whether there are any telltale signs of phenotypic dominance, i.e. patterns in the data that cannot be generated by additive models. In the current work, we have identified two. The first pattern involves Darwin’s Corollary, although it is not the observation of fitness asymmetry *per se* that implicates dominance. We have shown that log fitness asymmetry between cross directions tends to decline with genomic divergence under the additive model, but remains constant if there is variable phenotypic dominance (Fig. 5c–d). Note that the difference between predictions with and without dominance is relatively subtle, and so cannot be inferred directly from published results. For example, the relevant measure of asymmetry (eq. 19) uses log transformed fitness, relative to the maximum obtainable value, and this is difficult to obtain at low divergences – where parental fitness cannot serve as a proxy for the optimum – and at very high divergences – where zeros cannot be log transformed. The pattern can also be noisy (see Fig. S5e–f).

The second pattern that can only be explained by phenotypic dominance is somewhat simpler. Dominance and uniparental effects both allow for transgressive trait variation (i.e trait values that lie outside of the range of parental values; Stelkens and Seehausen, 2009), but only with dominance do the reciprocal cross directions produce transgression in the same phenotypic direction (Fig. 4a). The unique prediction of the dominance model is therefore a particular type of F1 fitness advantage. First, the advantage must apply to both cross directions. Second, it must not be due to the masking of deleterious recessives, nor to bounded hybrid advantage, such as found on an environmental gradient (with hybrids expressing an intermediate phenotype, beneficial in intermediate environments as in Wang et al., 1997); both latter sorts of heterosis are generated by the additive model (Simon et al., 2018; Schneemann et al., 2020; Yamaguchi and Otto, 2020). To demonstrate the pattern of heterosis that is unique to dominance, the reciprocal F1 must outperform both parents in a novel environment, while both parents outperform the F1 in their shared original environment. Equivalently, the two “environments” might involve the presence or absence of novel stressors, to which parental lines are not normally exposed. Some possible examples of such heterosis include increased thermal tolerance in copepod F1 (Pereira et al., 2014), invasive hybrid tiger salamanders (Ryan et al., 2009; Cooper and Shaffer, 2021), and *Senecio* ecotype hybrids in a habitat foreign to both parents (Walter et al., 2020). Such data are the strongest evidence for the importance of phenotypic dominance for F1 hybrid fitness.

This paper has focused on the consequences of dominance for the F1, but of course later-generation recombinant hybrids will experience some effects of dominance as well. While the telltale patterns in the F1 apply whenever dominance effects are present, the key question for later-generation hybrids is more quantitative: how do the deleterious effects of dominance effects compare to the costs of segregation variance, i.e. to the lack of coadaptation between the additive effects? We have denoted as *V*_*δ*_ the relative sizes of these effects, and shown that under some conditions, *V*_*δ*_ will equal the variance in the dominance coefficients of the fixed effects. The importance of dominance effects for later generation hybrids therefore depends on the size of *V*_*δ*_.

There are a number of observations that do provide upper bounds for *V*_*δ*_ in nature. The first is the ubiquity of Haldane’s Rule (Haldane, 1922; Coyne and Orr, 2004, Ch. 8; Schilthuizen et al., 2011). We have shown that Haldane’s Rule is predicted only when *V*_*δ*_ is smaller than the proportion of divergent loci that are biparentally inherited (eq. 18; Fig. 5a–b). Moreover, this bound applies to dominance effects at X-linked loci, which are probably larger than those on the autosomes (Charlesworth et al., 1987). Similar bounds also follow from direct observations of later-generation hybrids. For example, the model predicts that heterozygosity will be beneficial on average if *V*_δ_ < 1, and so this bound is supported by studies reporting selection for increased heterozygosity in recombinant hybrids (Lindtke et al., 2012; Simon et al., 2018; Thompson et al., 2022). Easier to observe, and so better supported, is hybrid breakdown—the fact that later-generation hybrids are very often less fit than the initial F1 (Muller, 1940; Vetukhiv, 1954; Fraïsse et al., 2016, Table S1). A simple generalization of eq. 18 (eq. 52) shows that hybrid breakdown between well adapted parental lines will occur only if:

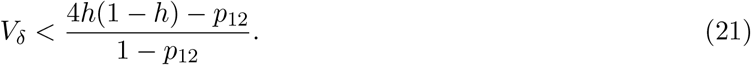

where *h* is the hybrid index (the proportion of divergent alleles that come from one the parental lines) and *p*_12_ is the interpopulation heterozygosity. It follows that F2 breakdown will occur whenever *V_δ_* < 1 (since 4*h*(1 − *h*) ≈ 1 for an F2). But backcross breakdown sets a stronger bound as it occurs only if *V*_*δ*_ < *p*_12_ (since 4*h*(1 − *h*) = *p*_12_(2 − *p*_12_) for any backcross), and *p*_12_ = 1/2 on average for the first backcross.

Taken together, then, these bounds do not allow us to conclude that dominance effects will be negligible for any class of hybrid. However, they are at least consistent with a world where 0 < *V*_*δ*_ ≪ 1. In this case, simple additive phenotypic models would yield good predictions for later generation recombinant hybrids (Fraïsse et al., 2016; Simon et al., 2018), even when dominance qualitatively affects outcomes for the F1.

## Supplementary Figures

**Figure S1:**
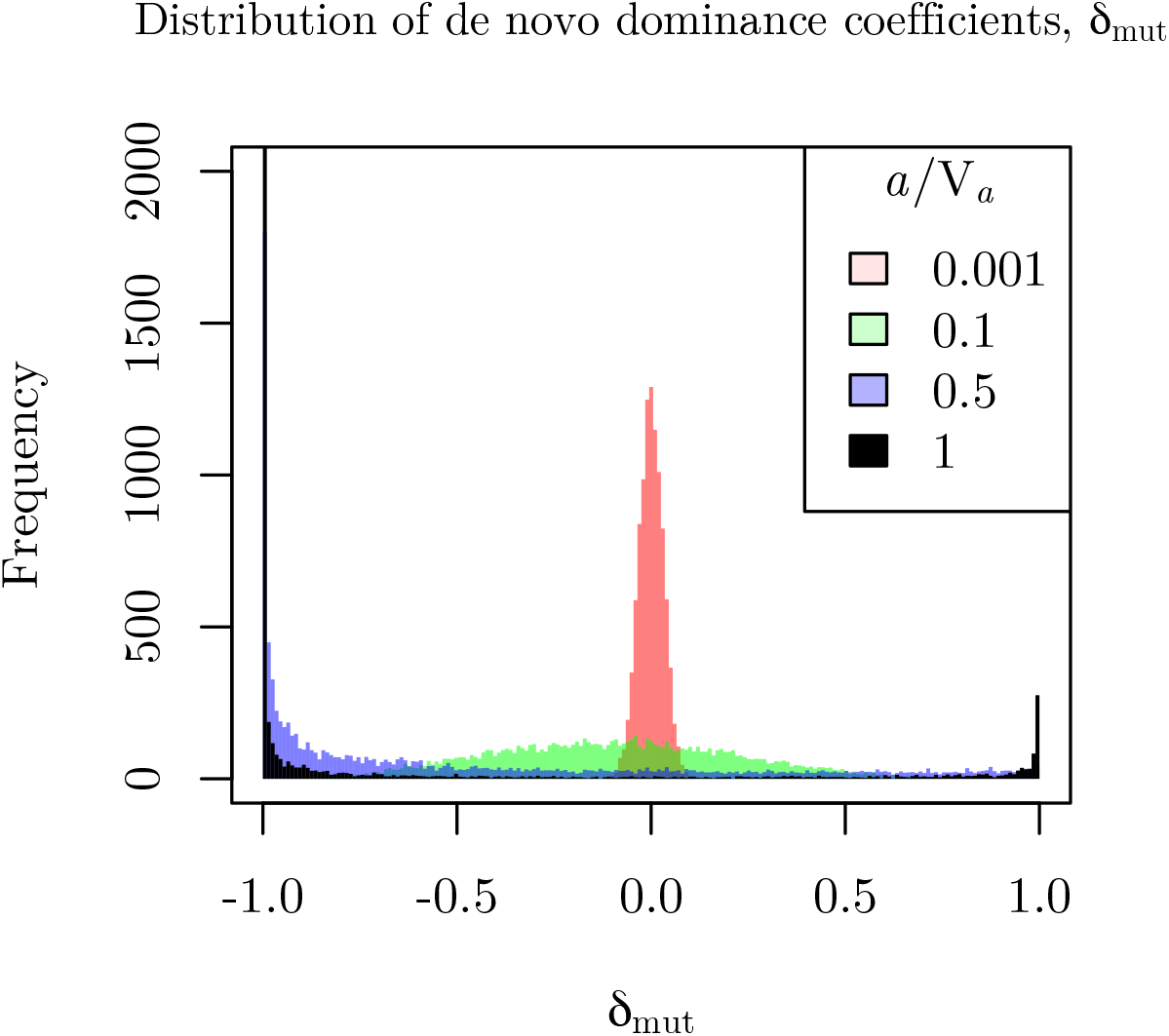
Size-dependent distribution of dominance coefficients. The distribution of dominance coefficients for new mutations as used in the simulations reported in Figures 1c, 4b and Figure S5. The heterozygous effect of a mutation on a given trait is *a*_mut_(1 + *δ*_mut_). Here, *a*_mut_ is the mutation’s additive effect, which is normally distributed with vanishing mean and standard deviation 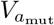 (eq. 59). The dominance coefficient, *δ*_mut_, is drawn from a beta distribution, whose parameters vary with the size of of the additive effect according to eqs. 60-61. The aim is to reflect empirical observations that lethal or highly deleterious mutations are often recessive, while small-effect mutations are more likely to be semi-dominant (Orr, 1991; Manna et al., 2011).

**Figure S2:**
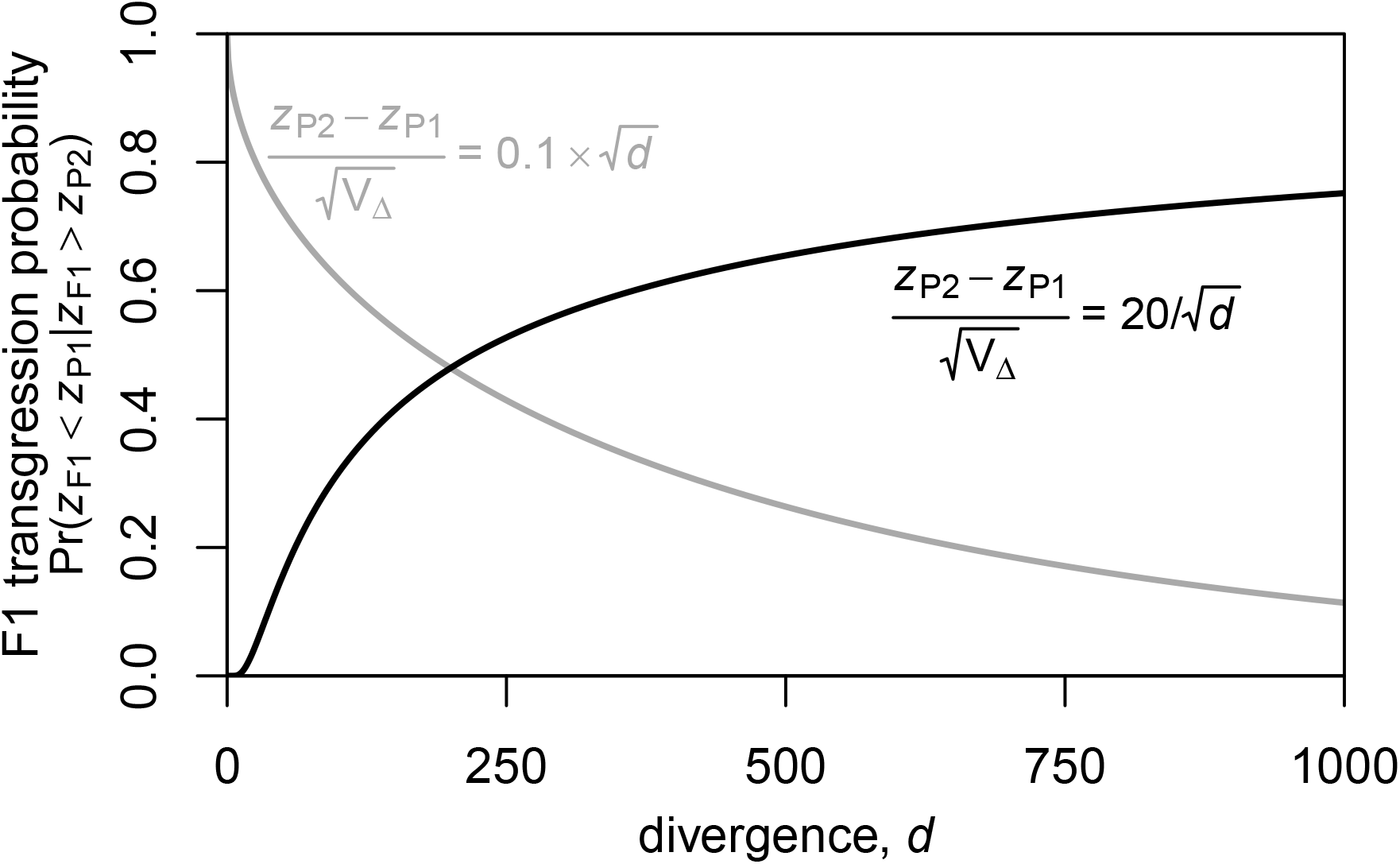
Transgressive F1 trait variation can increase or decrease with genetic divergence. Curves show the probability that the F1 trait value lies outside of the range of values in the two parental lines (eq. 47). The variance in the dominance coefficients, *V*_Δ_ increases linearly with genomic divergence, *d* (e.g. eqs. 6-7). If the divergence in the parental phenotypes also increases with *d* (e.g. due to directional selection) then the probability of transgressive variation decreases with *d* (grey curve). If the parental phenotypes remain roughly constant, e.g., due to effective stabilizing selection, then the probability of transgression increases (black curve). Both regimes have been observed in data (Stelkens and Seehausen, 2009).

**Figure S3:**
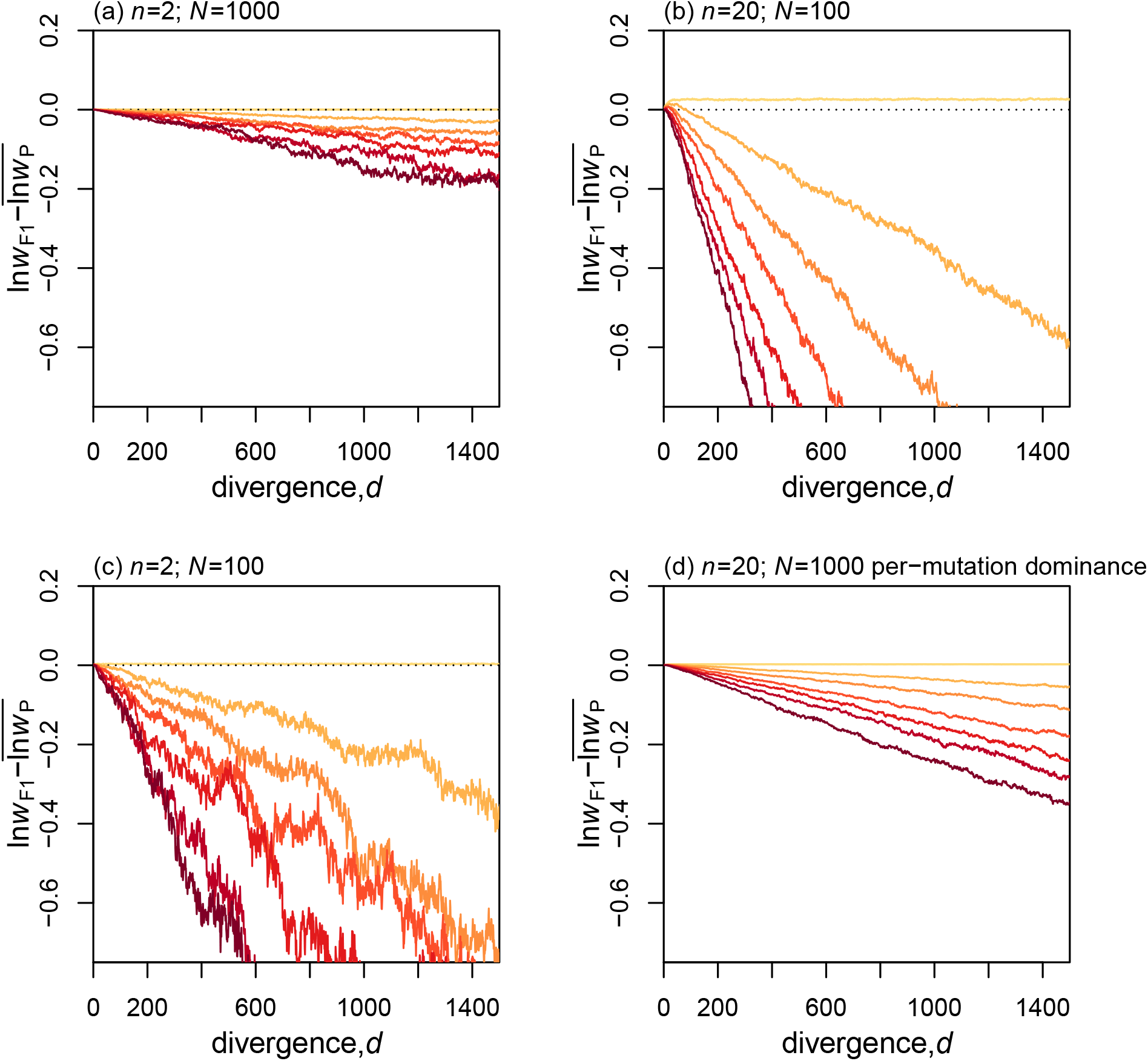
The F1 clock tick rate varies with various model parameters. **(a)-(c)**: Results replicate those shown in Figure 2b, except that the simulation parameters differed in terms of the number of traits (a), the population size (b), or both (c). In all cases, the slopes get steeper with 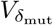, but the decrease is not always linear, showing that 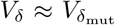 does not always hold. (d): Results also hold in simulations where each mutation had a single value of *δ*_mut_ applying to all *n* traits; such that heterozygous effects differed from the homozygous effects in their length but not in their direction. This shows that the effects of dominance reported here do not depend on the variation in direction.

**Figure S4:**
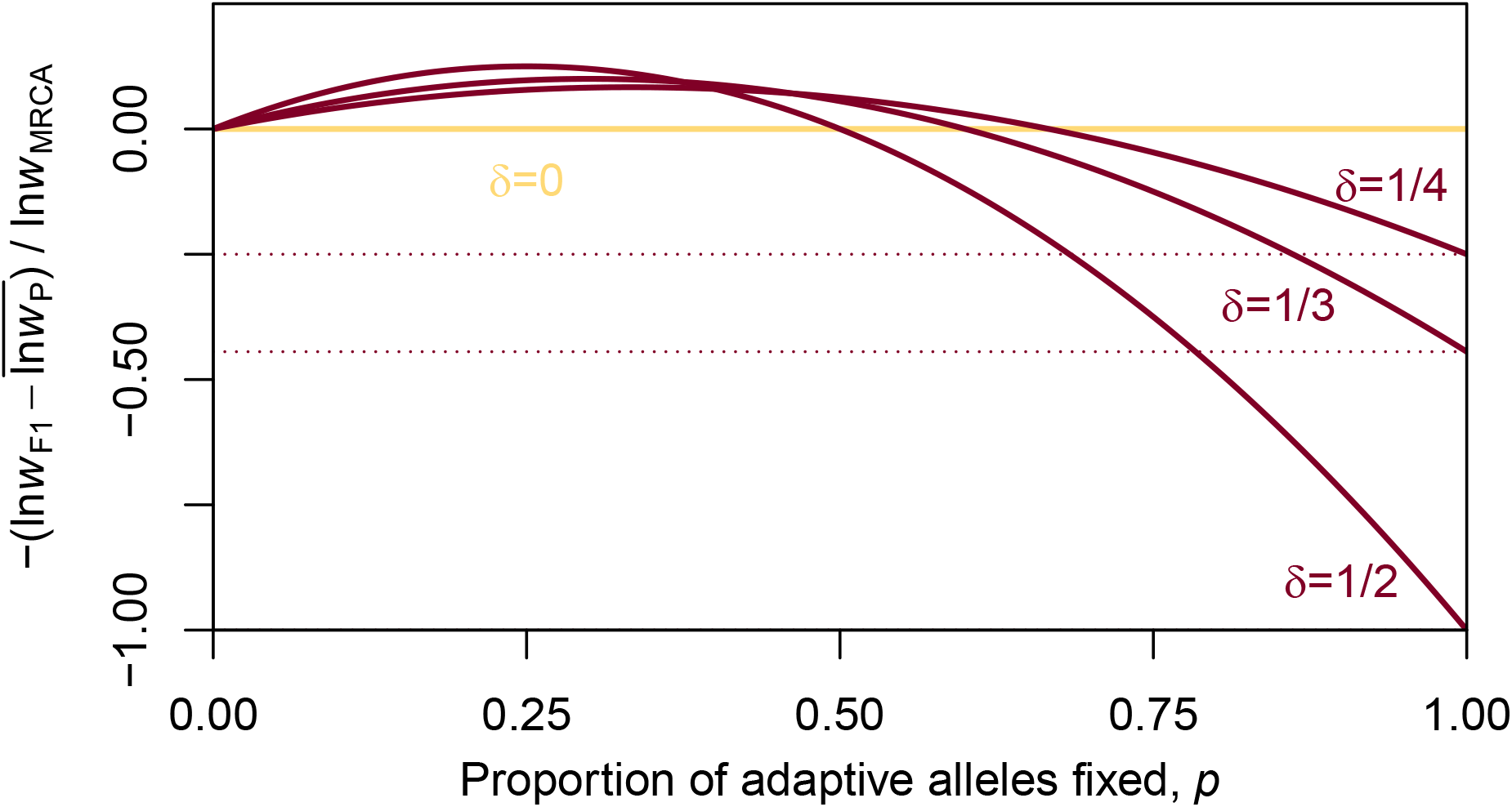
Directional dominance leads to a transient fitness increase and then a permanent decrease. Log F1 fitness relative to current and pre-adaptation parental fitness when parental lines are approaching a new optimum through the fixation of identical substitutions with equally sized additive effects and dominance effects (see eq. 37). The size of the dominance coefficients is denoted *δ*. The yellow line *δ* = 0 represents additivity, and the dark red curves represent dominance coefficients of different magnitudes. With dominance, F1 fitness first exceeds parental fitness due to the directional dominance overshoot, but as parental lines get closer to the new optimum this overshoot becomes deleterious and F1 fitness declines. The horizontal dotted lines give the predicted overshoot when the optimum has been reached.

**Figure S5:**
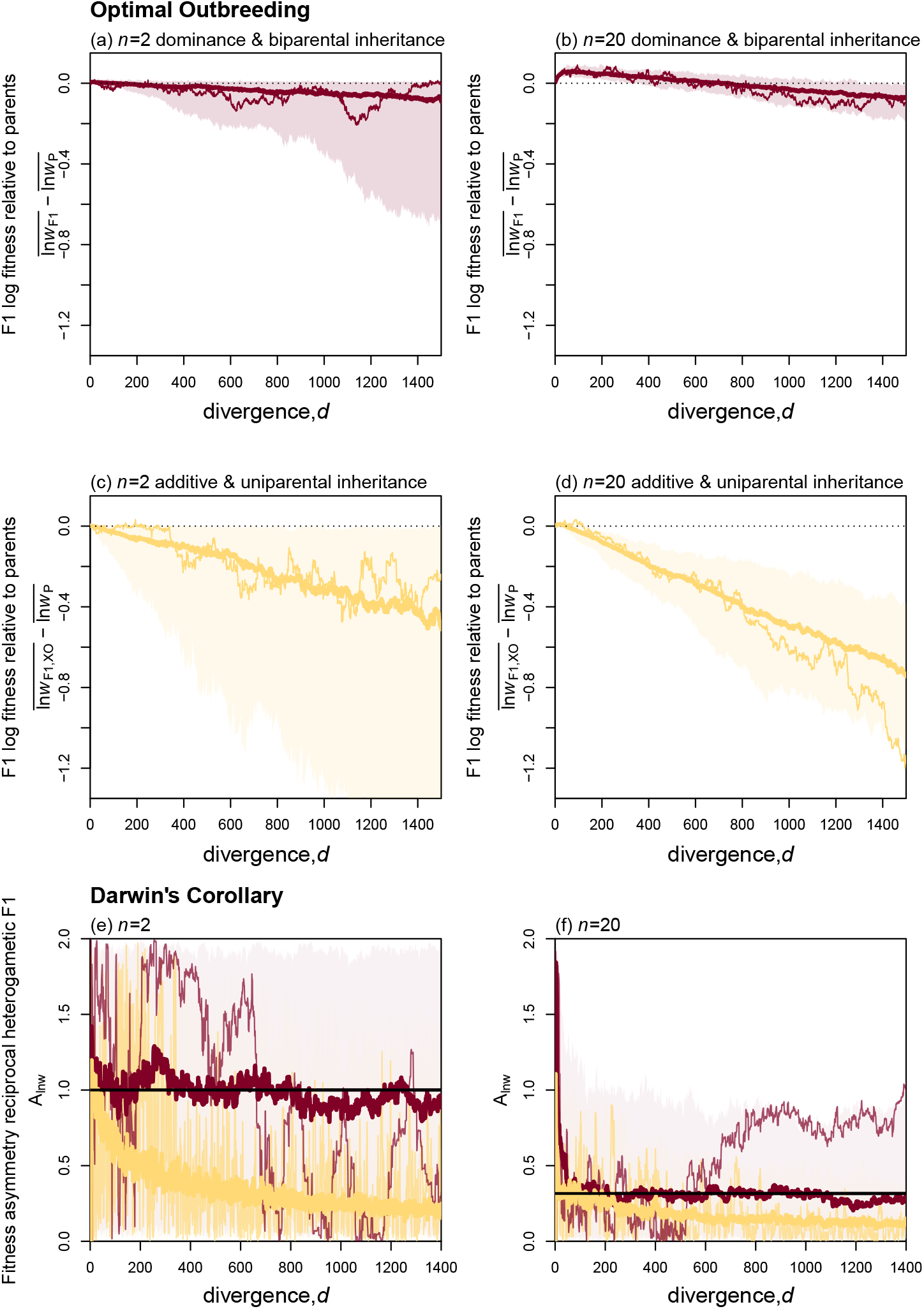
Major patterns of F1 fitness for simulated F1 hybrids using the dominance function illustrated in Fig. S1. Simulation results are reported comparing phenotypic additivity (yellow curves) to phenotypic dominance, using the dominance function shown in Fig. S1, where mutations of larger effect had more extreme levels of dominance (dark red curves). Results correspond to **(a)**-**(b)**: Fig. 1c; **(c)**-**(d)**: Fig. 4b; **(d)**-**(e)**: Fig. 5c–d. In all cases, the average over 100 replicates (thicker solid lines) is compared to a single randomly chosen replicate (thinner lines), to illustrate the stochasticity. Heterogametic F1 in (c)–(f) used 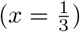.

**Figure S6:**
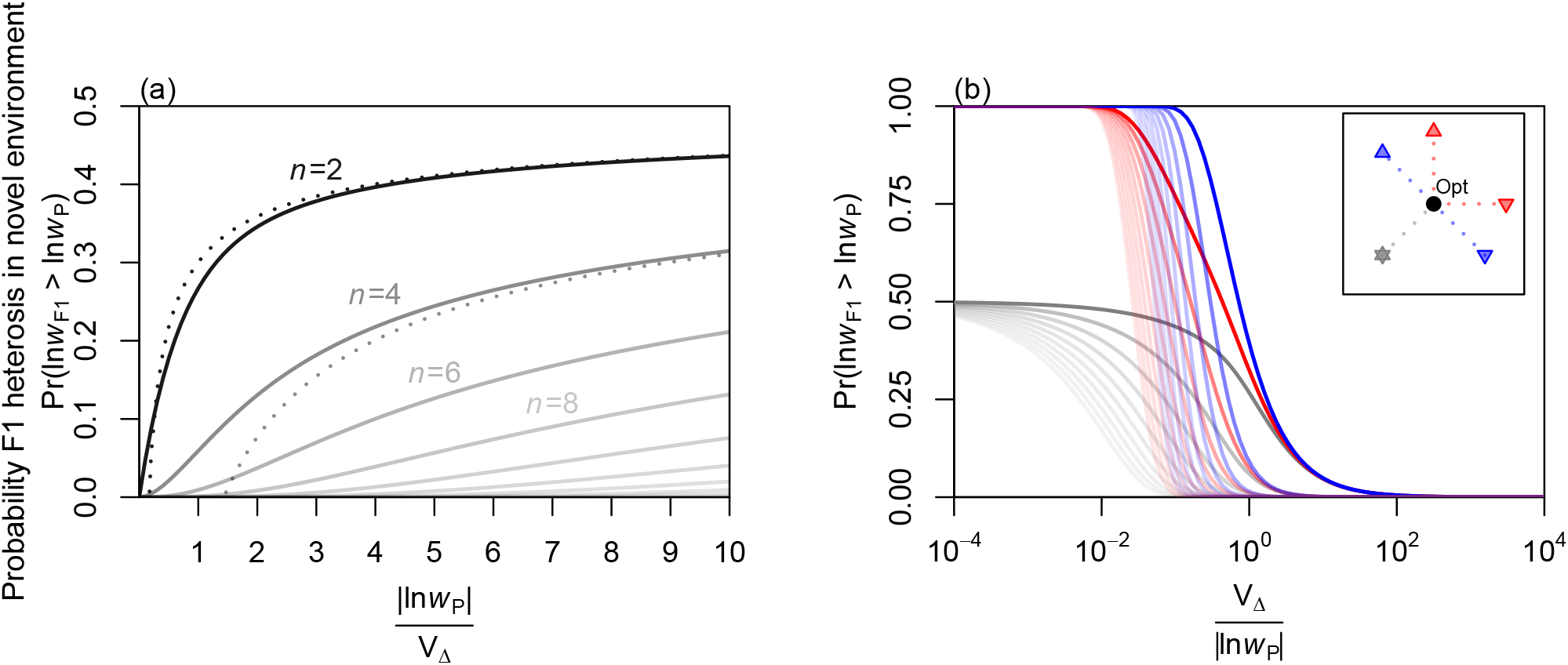
The probability of F1 heterosis in a novel environment depends on the number of traits, and the parental phenotypes. Panel **(a)** and gray lines in **(b)** represent the scenario from Figure 3a, when the two parental lines have similar and equally maladapted phenotypes. **(a)**: a replication of Figure 3b, but without log transformation on the x-axis, highlighting the decline in the probability of heterosis with *n*. **(b)**: the probability of heterosis increases if the parents are maladapted on different traits (red curves), or in opposite phenotypic directions (blue curves). The positions of the parents relative to the optimum are shown in the inset. Colour intensity indicates the value of *n* which ranges from 2 (bright lines) to 20 (faint lines), and in all cases, the probability of heterosis declines steeply with *n* and with *V*_Δ_/| ln *w*_P_|. The functions plotted are described in Appendix 1.

**Figure S7:**
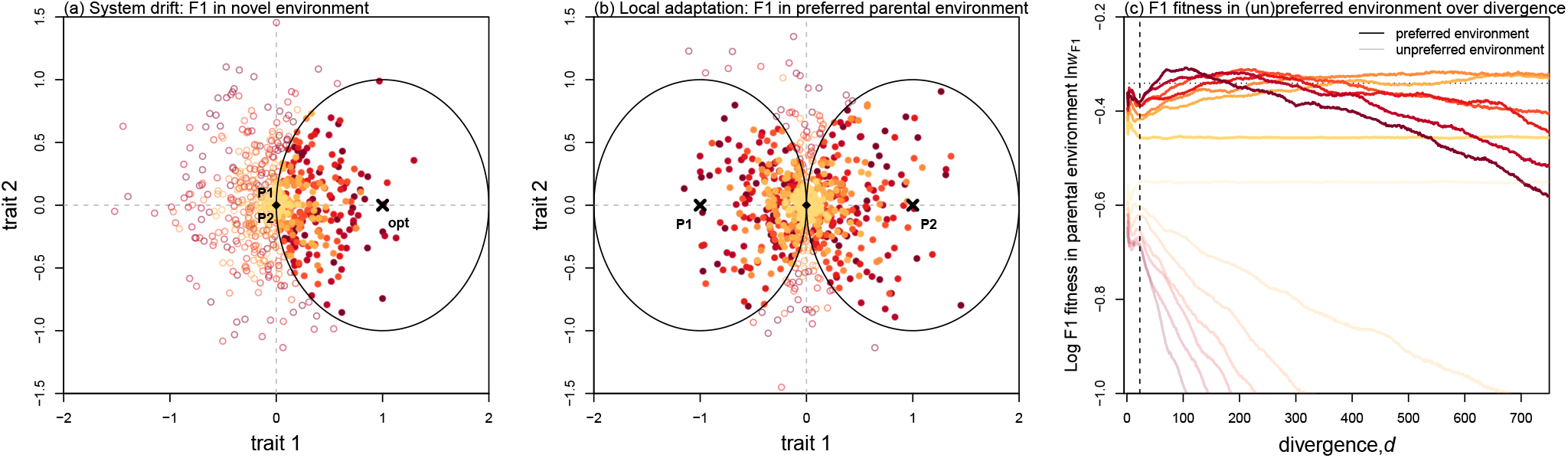
Lucky beneficial effects of dominance in novel environments, with variable levels of mutational dominance. **(a)**–**(b)**: Simulated F1 hybrids after 50 divergent substitutions with different levels of variation in dominance show “Lucky” beneficial effects in a new environment. **(c)**: Simulations of the expected log fitness of the F1 between locally adapted parents, scored in the preferred parental habitat, to which the F1 is better adapted (solid lines) or the unpreferred parental habitat, to which it is less adapted (faint lines). These results match Figure 3d, but include results for all seven different levels of variance among the dominance coefficients of the new mutations, 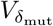. Colours for the 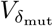 values match those in Figure 2, with darker colours indicating higher mutational variance.

**Figure S8:**
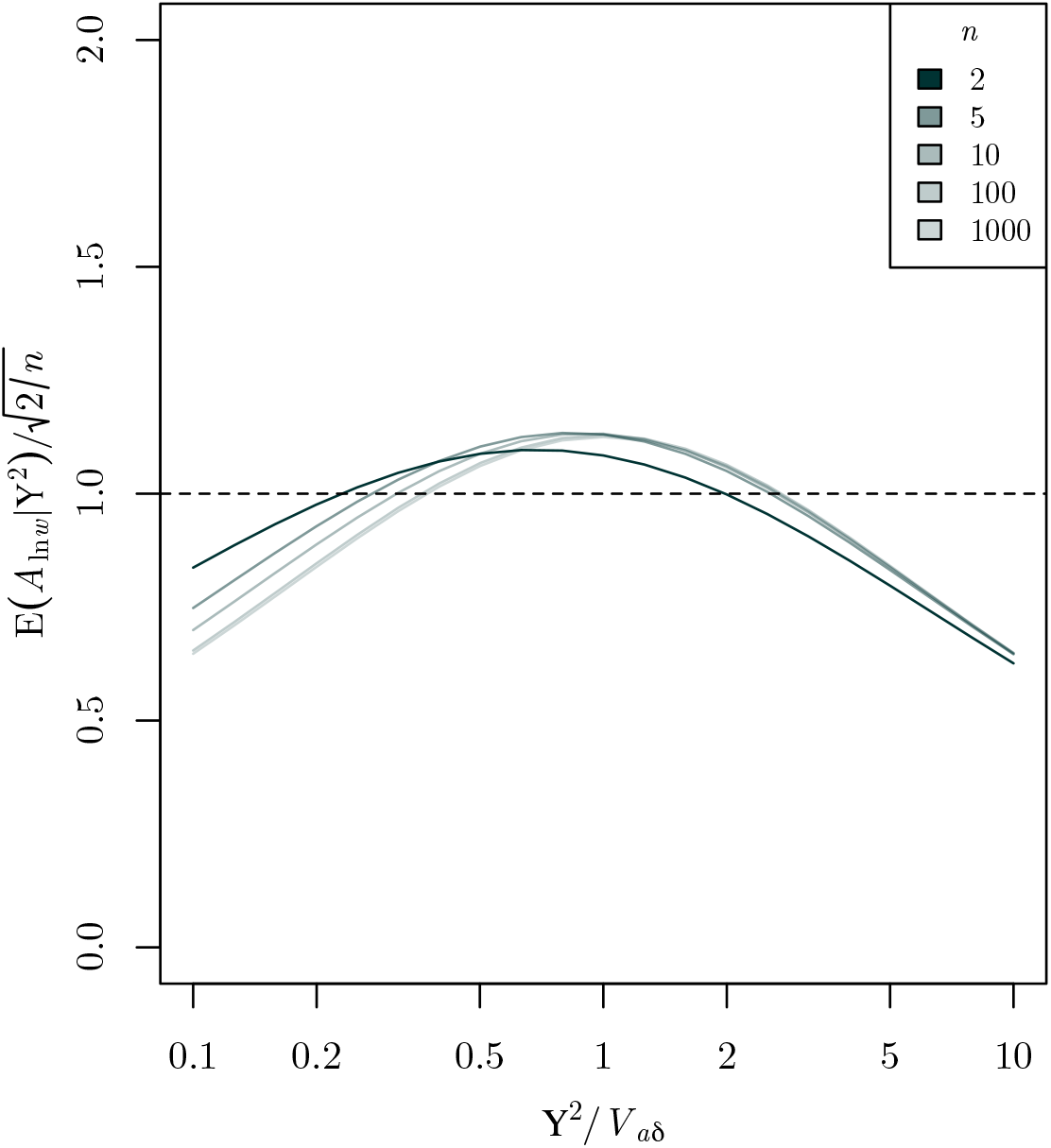
Derivation of the expected levels of log fitness asymmetry between cross directions with variable phenotypic dominance. The expected value of eq. 55 is shown for various values of *n*, namely (dark-to-light) *n* = 2, 5, 10, 100 and 1000. The expectation (evaluated numerically) is close to the value of the crude approximation 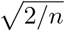 (eq. 20), unless the quantity Υ^2^/*V*_*aδ*_ deviates greatly from unity. This quantity compares the deviations due to dominance, and the deviations due to lack of autosomal co-adaptation with the absent paternal X chromosome (see Figure 4a), and depends mainly on the variance in the dominance effects and the proportion of uniparentally inherited loci (eq. 57).

## Appendix 1: Derivations

In this Appendix, we derive the analytical results in the main text. The relevant main text equations are listed next to each section.

## 1 F1 fitness with biparental inheritance

Many results in the main text are given in terms of the total dominance deviations on each trait, that is, the Δ*_i_* as defined in eq. 4. The strongest assumptions that we use are the normality and isotropy of the Δ*_i_* (i.e. that all Δ*_i_* have equal variances). However, neither assumption is required for all results, and the assumption of isotropy is made largely for simplicity.

## 1.1 Expected log fitness (eqs. 6–11)

s

To calculate the expected log fitness of the F1, we begin by expanding the brackets in eq. 5:

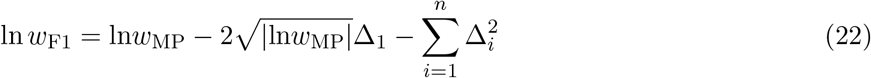

Then we take expectations over the possible values of the dominance deviations, to account for their relative independence of the form of selection acting on the parental lines.

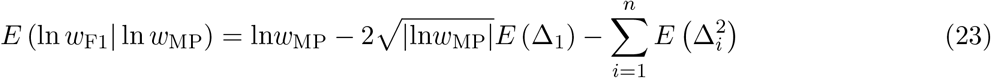

Equations 9–10 follow directly, if we focus only on trait 1.

## 1.1.1 Stabilizing selection (eqs. 6–8)

Next, if there is no directional dominance, i.e., if all *E*(Δ*_i_*) = 0, then:

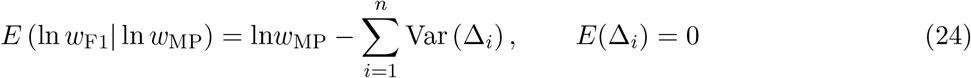

and so eq. 6 follows directly if we use the notation:

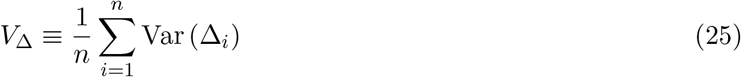

For eq. 7, we use equivalent notation to refer to the effects of the individual substitutions, which under stabilizing selection, we can treat as increments in a random walk, with variance

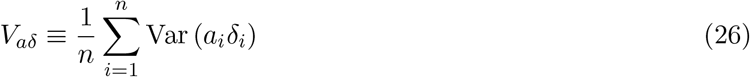

where *a*_*i*_*δ*_*i*_ is a randomly chosen dominance effect on trait *i*, and the expectations are over realizations of the evolutionary process. We can also define an equivalent quantity for the additive effects.

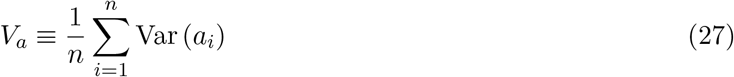

where *a*_*i*_ is a randomly chosen additive effect on trait *i*. Equation 8 now holds generally if we define the equivalent quantity for dominance coefficients, *V*_*δ*_ as the ratio of the two quantities above

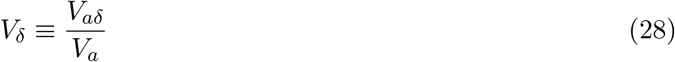

such that it indicates the size of the dominance effect relative to the additive effect of substitutions. *V*_*δ*_ is not generally equivalent to the variance in dominance coefficients, but it is equivalent in the special case of isotropy, and no covariance between the squares of the additive and dominance effects. To see this, we can also write *V*_*aδ*_ in terms of moments of additive effects and dominance coefficients, where similar to the above we use *δ*_*i*_ to refer to the dominance coefficient on trait *i* of a randomly chosen substitution. With exchangeable parental lines, and recalling that the effects of a substitution are defined with respect to P1, irrespective of whether they are ancestral or derived, it follows that *E* (*a*_*i*_) = *E* (*δ*_*i*_) = 0. Looking at the other moments, we find

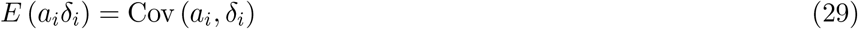

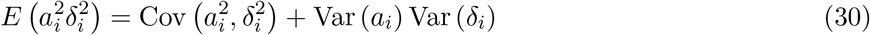

Now we can write *V*_*aδ*_, as

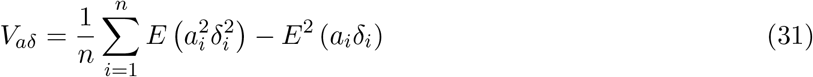

without directional dominance Cov (*a*_*i*_, *δ*_*i*_) = 0, and if we also assume that larger substitutions have no tendency to have more extreme dominance coefficients, then Cov 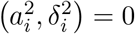, and we get

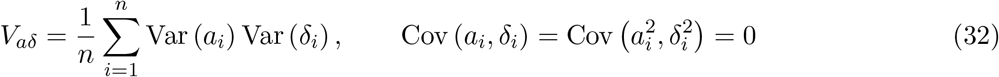

Under isotropy, the distribution of substitutions is the same across traits, so that *V*_*a*_ = Var (*a*_*i*_) and *V*_*δ*_ = Var (*δ_i_*) are both constant. Thus, with these additional assumptions we can again obtain eq. 8, but with a more intuitive intepretation of *V*_*δ*_. Note, however, that data do not support the assumption that 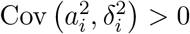 (Orr, 1991; Manna et al., 2011).

## 1.1.2 Directional selection (eqs. 9–11)

Under directional selection on trait 1, we expect additive effects in the direction of the optimum to be more dominant. Therefore, for these adaptive substitutions (denoted by an asterisk *), 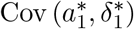 will not vanish. However, we can obtain results in a simplified case in which the initial bout of adaptation took place in both parental lineages, taking them both from an old optimum to a new optimum, and where these optima differed only on trait 1. We will assume that the adaptation took place in exactly *d*_dir_ substitutions, and that these substitutions affected only trait 1. We will further assume that their additive effects and dominance coefficients covary in sign, but not in size, such that 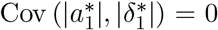. In other words, large adaptive substitutions are no more likely to be dominant or recessive than small adaptive substitutions, but substitutions pointing from the MRCA towards the optimum (adaptive substitutions that are derived in P2) have 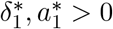 whereas substitutions pointing towards the MRCA and away from the optimum (adaptive substitutions derived in P1) have 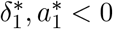.

Taken together, the chain of adaptive substitutions runs from the new optimum to the old optimum and then back, and so its total length is 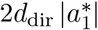, and we have 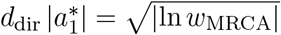, where *w*_MRCA_ is the fitness of ancestral genotype (adapted to the old optimum), in the environment characterized by the new optimum. With these assumptions, we can now derive the moments of Δ_1_ after both the adaptation, and a further period of *d* − *d*_dir_ substitutions, accrued under stabilizing selection:

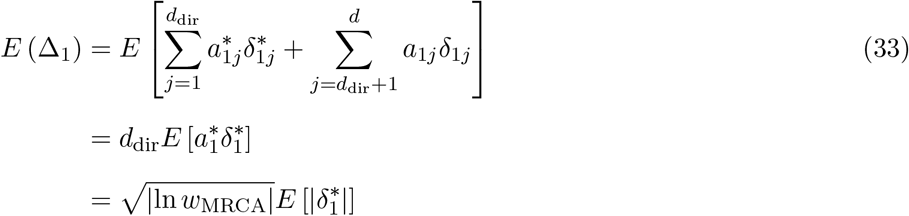

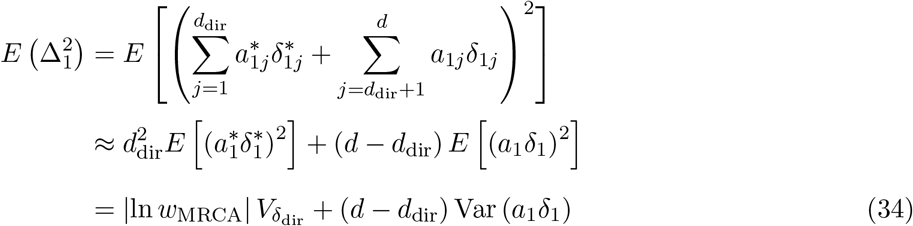

where

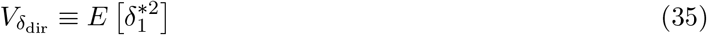

Here, we used the random-walk assumptions, described above, for the substitutions accrued after the adaptation. Equation 34 also uses the stronger assumption that covariance between the dominance effects of the adaptive substitutions can also be ignored. After making these assumptions, eq. 11 in the main text follows directly.

We can also give a more complete result that applies during the ongoing adaptation to the new optimum. During the adaptation phase i.e. before the parental lines have reached the optimum, we use 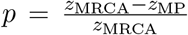 to denote the proportion of the distance to the new optimum the parental lines have already travelled. Considering only the trait under directional selection, we find

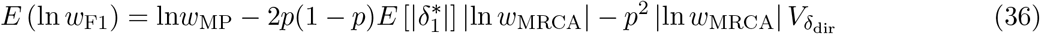

For the overshoot of the F1, we find

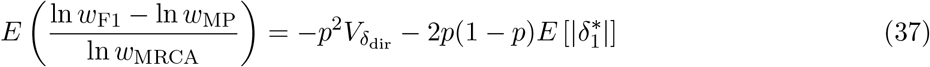

These expressions are plotted in Figure S4 for substitutions with dominance coefficients of identical size |*δ*|.

## 1.2 Variance in F1 log fitness (eq. 12)

To calculate eq. 12 we also require the variance in log F1 fitness. Its behaviour is easiest to see if we assume normality and independence of the Δ*_i_*. In this case, we find:

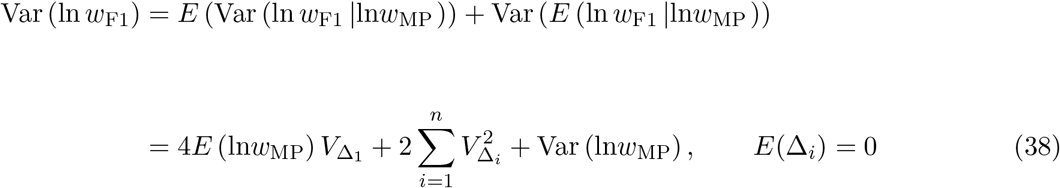

If the midparent remains well adapted, but the dominance deviations are large, then the second term will come to dominate. Because *V*_Δ_ grows with the divergence, this will typically hold at large divergences, when *d* ≫ 1, and so

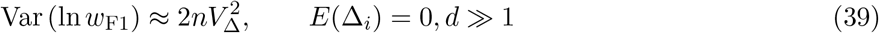

Using the same limit in eq. 6 yields eq. 12.

## 1.3 F1 fitness in preferred parental environment (eqs. 14, 15)

When the parental lines are adapted to two different optima, then the F1 will tend to be fitter in one of the two parental environments (i.e. its “preferred” environment). In that environment, we can write its log fitness using a small modification of eq. 22:

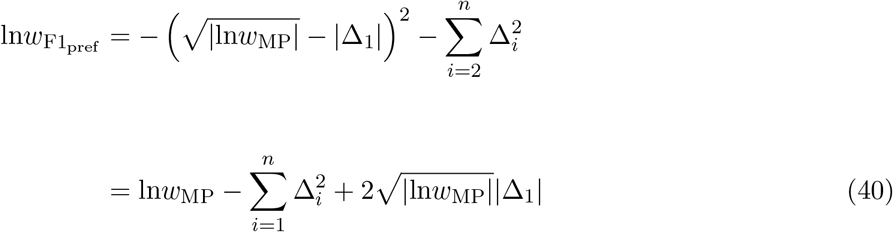

If we assume that the parental lines are each optimal in one of the habitats, then distance of midparent from either optimum is half of the distance between the optima, such that ln *w*_MP_ = ln *w*_P_/4, where ln *w*_P_ is the log fitness of the maladapted parental line in any given habitat. Next, we assume that Δ_1_ is normal, with a vanishing mean, so that 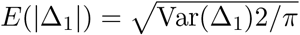. Equation 14 then follows directly if we assume equal variances for the dominance deviations on all traits, so that Var(Δ_1_) = *V*_Δ_.

To derive equation 15, we differentiate eq. 14 by *V*_Δ_, and set to zero. This yields the value of *V*_Δ_ that maximises the expected log fitness. Substituting that value into eq. 14 yields eq. 15.

As with eq. 39, we can also derive the variance 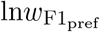 in using the assumptions of normality, independence and vanishing means for the Δ*_i_*:

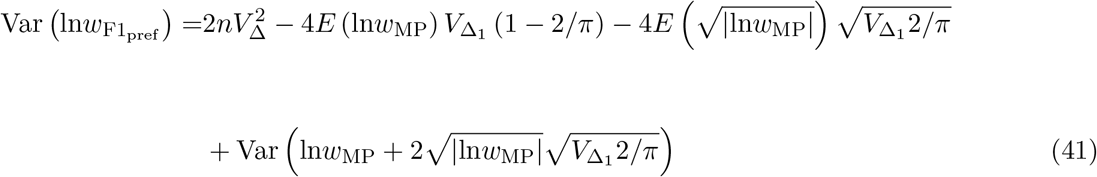

At large divergences, the first term will come to dominate, supporting the assertion in the main text that eq. 12 still holds for 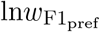.

## 1.4 Probability of lucky heterosis (eq. 13)

Now let us consider the F1 between phenotypically identical parental lines, that are both maladapted. To calculate the probability of their F1 being heterotic (Fig 3b), we first note that, with isotropy, such that all 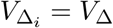, then, from eq. 5, the quantity | ln *w*_F1_|*/V*_Δ_ is the sum of *n* squared normal variables, one with a non-zero mean of 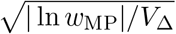. As such, this quantity will follow a non-central chi-squared distribution, with *n* degrees of freedom, and non-centrality parameter given by | ln *w*_MP_|*/V*_Δ_ = | ln *w*_P_|*/V*_Δ_. Here, we have used the fact that the midparent will be identical to the two (identical) parental phenotypes. We can write the cumulative distribution function of this variable as

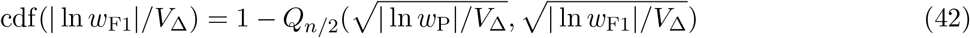

where we have used the Marcum Q function (Marcum, 1950). The F1 will be heterotic if it is closer to the optimum than the midparent and so

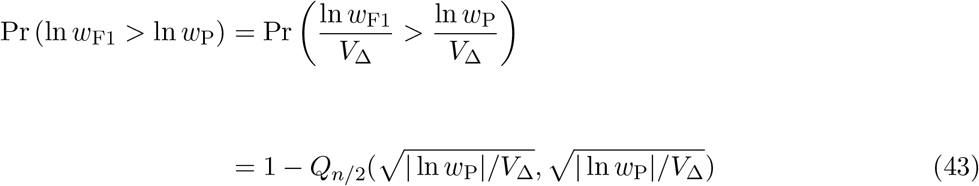

This is the exact result plotted in Figure 3b. This result can also be written in terms of modified Bessel functions of the first kind, as follows:

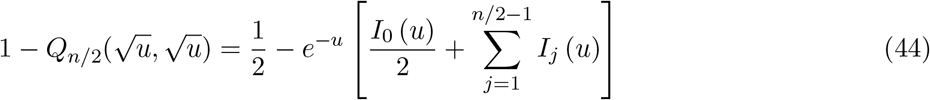

(Helstrom, 1995). Approximating these functions as 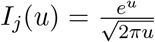 (see, e.g. Guo et al., 2001), yields eq. 13 of the main text.

Figure S6 shows similar results when the parental lines are equally maladapted, but no longer phenotypically identical. To derive these results, we simply change the non-centrality parameter (i.e., the first argument of the Q function in eq. 43), which represents the position of the midparent. When the parents are maladapted in opposite directions, then the midparent matches the optimum, such that this argument is 0 (blue points and lines in Fig. S6). When the parents are maladapted in orthogonal directions (e.g., on different phenotypic traits), then this argument is 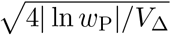 (red points and lines in Fig. S6).

## 1.5 Transgressive segregation in the F1

The procedure used in the previous section can also be used to calculate the probability of transgressive variation in a single trait: i.e., an F1 trait value that lies outside of the range of values among the parental lines (Stelkens and Seehausen, 2009). If there is no directional dominance, then the mean F1 trait value will be the midparental value, and its variance will be equal to the variance of the dominance deviations.

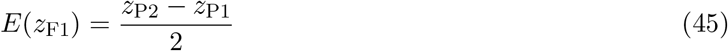

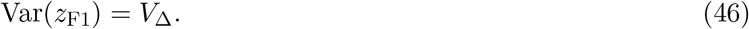

where we have ordered the parents by their trait value. Now, assuming normality, the probability of the F1 trait value lying outside of the parental range is

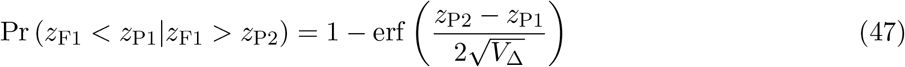

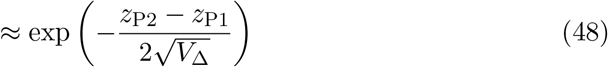

Because *V*_Δ_ grows linearly with *d*, the probability of transgressive variation will increase with *d* if the divergence between the parental phenotypes also increases with *d* (at least as fast as 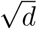. If, by contrast, the difference between the parental traits remains static, then the probability of transgressive variation will decrease with *d*. This is illustrated in Figure S2. Both regimes were observed by Stelkens and Seehausen (2009).

## 2 Uniparental inheritance and recombinant hybrids

In this section, we present results for a more general type of hybrid, in which some loci are either hemizygous or homozygous. To do this, we will assume that the the parental lines are under stabilizing selection, as described above, such that the dominance deviations undergo a random walk in phenotypic space. We also use existing results for recombinant hybrids, conditional on the F1 fitness (Simon et al., 2018; Schneemann et al., 2020; De Sanctis et al., 2022 in prep.).

## 2.1 Expected log fitness and bounds on *V*_*δ*_ (eqs. 16–18 and 21)

An arbitrary hybrid can be characterized by its hybrid index, 0 ≤ *h* ≤ 1 (i.e. the proportion of divergent alleles from one of the parental lines) and by its heterozygosity, 0 ≤ *p*_12_ ≤ 1 (the proportion of divergent loci carrying one allele from each parental line). Let us first begin by defining *A*_*i*_ as the analogue to Δ*_i_* for the additive effects.

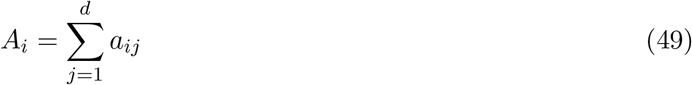

With this notation, the midparental log fitness can be written as

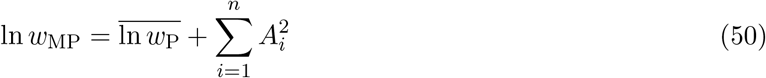

It then follows from eq. 3, that when the populations are maladapted from their optimum in random directions, such that 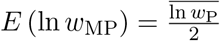, then

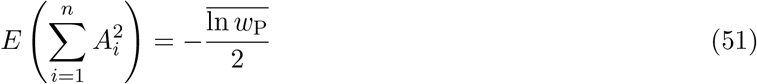

Now, using the assumptions discussed above for stabilizing selection, we can combine eq. 51 with the published results, to yield the expected log fitness of an arbitrary hybrid.

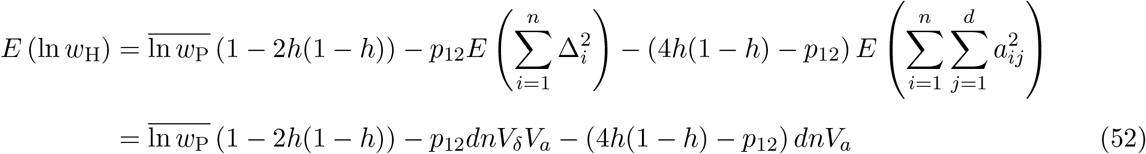

Note that eq. 52 contains three terms, the second capturing the deleterious effects of dominance, and the third capturing the deleterious effects of lack of coadaptation among the additive effects. Equation 52 also reduces to results in the main text for a fully heterozygous F1, for which *h* = 1/2 and *p*_12_ = 1. Equation 21 follows from comparing this F1 to the arbitrary hybrid, when parents are well adapted 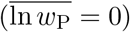. Equations 16–18 also follow simply. For an XO F1 hybrid, all of the heterozygosity is found on the autosomes, such that *p*_12_ = 1 − *x* while the hybrid index is *h* = (1 ± *x*)/2, depending on which parental line contributed the X. It follows, therefore, that 2*h*(1 − *h*) = (1 − *x*^2^)/2, and 4*h*(1 − *h*) − *p*_12_ = *x*(1 − *x*).

## 2.2 Darwin’s corollary (eq. 20)

For the asymmetry of log fitness between the cross directions, we will begin by assuming that the parental phenotypes – and therefore the midparental phenotype – are optimal. This means that the reciprocal XO F1, carrying different X chromosomes, will deviate in equal and opposite extents from the optimum. We will denote these deviations as 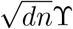, reflecting their growth with the number of traits, and with divergence (e.g. eq. 52). The dominance deviations, which come from their shared autosomes, will be in consistent directions. As such, we can orientate the trait axes so that these deviations fall on trait 1. This means that the difference in their log fitnesses can be written as:

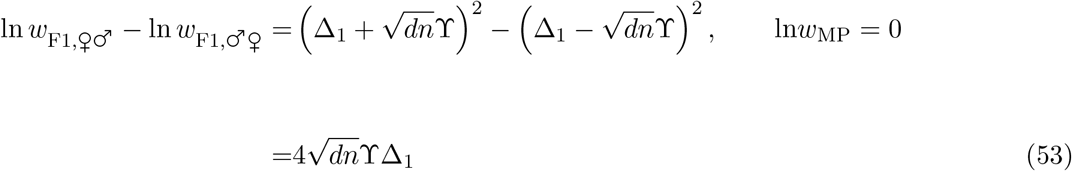

while the mean across the cross directions yields:

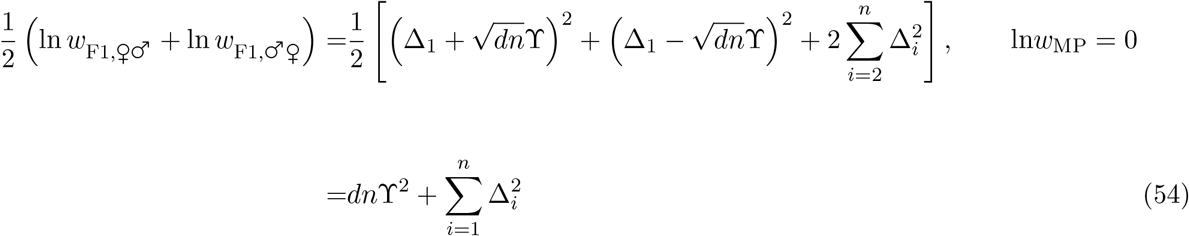

We now treat the dominance deviations as normal variables, such that 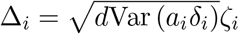, where *ζ_i_* is a standard normal random variable, with vanishing mean and unit variance. The measure of asymmetry is therefore

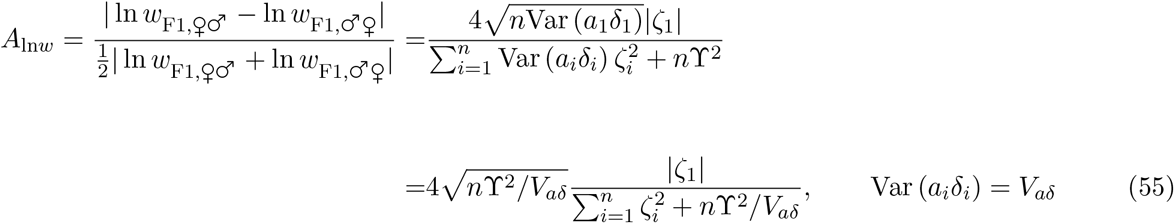

So this quantity is not expected to change in any consistent way with the genetic divergence, *d*. We have not been able to evaluate this expression analytically, but numerical integration shown in Figure S8, shows that, to a rough approximation

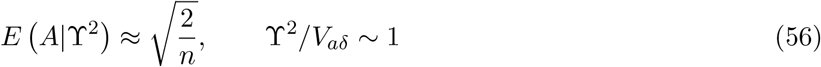

which agrees with the simulation results reported in the main text (Figure 5c–d) and in Figure S5e–f. We can also anticipate the typical value of Υ^2^/*V*_*aδ*_, using eq. 16. This shows that

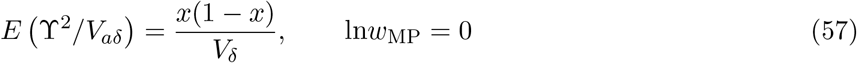

suggesting that Υ^2^/*V*_*aδ*_ ~ 1 will often hold for plausible parameter values.

## Appendix 2: Simulation procedure

To complement our analytical predictions, we used individual-based simulations of evolutionary divergence under Fisher’s geometric model. The simulation procedures were closely based on those reported in Schneemann et al. (2020), and for compatibility with earlier work, we included a constant into the log fitness function, such that:

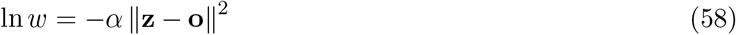

All simulations used *α* = 1/2. In all cases, we simulated pairs of allopatric diploid populations, which were initially identical with no segregating variation. The ancestral population was phenotypically optimal for all simulations except for those presented in Figures 2e–f (modelling directional selection) and 3d (modelling locally adapted parents), where the ancestor was placed at unit distance from the optimum, such that the ancestral population had a relative fitness of *e*^−*α*^ ≈ 61%, and the simulations began with a bout of directional selection towards the optimum. In all cases, the diverging populations were of constant size *N* individuals (2*N* chromosomes), and contained simultaneous hermaphrodites with discrete non-overlapping generations.

To produce the next generation, 2*N* prospective parents were sampled with replacement, with probabilities proportional to their fitness. Gametes were produced with free recombination among all loci, and assuming infinite sites, and subject to mutation. The total number of mutations was drawn from a Poisson distribution with mean 2*NU* and distributed randomly across the 2*N* gametes. There was no back mutation. The mutation rate *U* was chosen such that multiple mutations generally segregated together (*NU* ≥ 1). The additive effects of each mutation on each each trait, were drawn independently from a normal distribution, with vanishing means and variance denoted 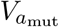. We set this mutational variance at:

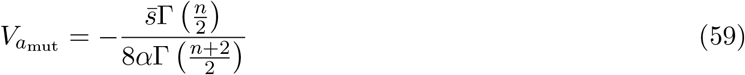

Because the homozygous effect of a mutation was twice its additive effect, eq. 59 ensures that the mean selection coefficient acting on a homozygous mutation occurring in an optimally fit individual was approximately 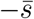, irrespective of the number of traits under selection, *n*.

The heterozygous effect of a mutation was equal to its additive effect, *a*_mut_, multiplied by (1 + *δ*_mut_), where *δ*_mut_ is its dominance coefficient. For simulations under phenotypic additivity, we simply set *δ*_mut_ = 0 for all mutations. With variable phenotypic dominance, the *δ*_mut_ were drawn from shifted beta distributions, which we bounded at −1 (for a wholly recessive mutation) and 1 (for a wholly dominant mutation).

For the simulations reported in Figures 2, 3d, 5, and S3, we used the same shifted beta distribution for all mutations, regardless of their additive effects. To do this, we set set *E*(*δ*_mut_) = 0 and 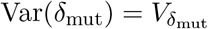. These distributions are shown in Figure 2a. Generating dominance coefficients in this way allowed us to adjust the amount of variance in dominance coefficients of new mutations 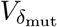, which was reflected in *V*_*δ*_.

For the simulations reported in Figures 1c, 4b and S5, we generated the *δ*_mut_ values to reflect some of the known properties of large and small effect mutations. Specifically, small effect mutations were close to additive with little variance, whereas large effect mutations were predominantly recessive but with a higher variance, such that some would be dominant (Orr, 1991; Manna et al., 2011). This was achieved by sampling the dominance multipliers from a beta distribution such that

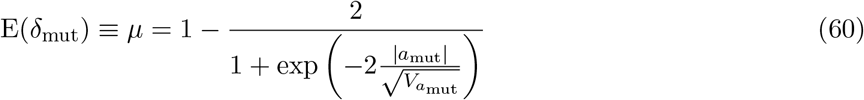

and

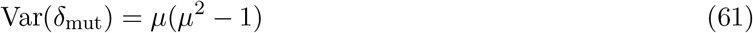

The distributions generated by eqs. 60–61 are illustrated in Figure S1.

For each set of parameters, we simulated 200 independent populations that each evolved until they fixed 1000 substitutions. We then randomly paired populations to obtain 100 pairs for the formation of F1 hybrids. We considered the substitutions that had fixed in the number of generations it took for the first population of the pair to reach 1000 substitutions. To produce the F1, for standard biparental inheritance (and also for homogametic hybrids), we took the heterozygous effect of all substitutions. For our heterogametic hybrids, which were our example case of uniparental inheritance, we assigned a random proportion *x* of substitutions to be on the X chromosome, and considered their homozygous rather than their heterozygous effects. This is equivalent to a form of dosage compensation, involving up-regulation on the X (see main text). Note that these X-linked substitutions were chosen after simulating the divergence. This meant that all sites were biparentally inherited and expressed during the divergence process. This guaranteed that, for our simulations, the variance in dominance coefficients of fixed mutations did not differ between the X and autosomes, with both equal to the mutational input, such that 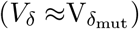 This made it easier to illustrate the conditions for Haldane’s Rule, eq. 18 (see Figure 5), but it is unlikely to hold in nature (Charlesworth et al., 1987, see also Discussion).

